# A transcytotic actin shift polarizes vesicle trajectories and partitions apicobasal epithelial membrane domains

**DOI:** 10.1101/2022.02.06.479326

**Authors:** Gholamali Jafari, Liakot A. Khan, Hongjie Zhang, Edward Membreno, Siyang Yan, Verena Gobel

## Abstract

In prevailing epithelial polarity models, membrane-based polarity cues such as the partitioning-defective PARs specify the positions and identities of apicobasal membrane domains. Recent findings suggest, however, that vesicle-associated polarity cues specify membrane polarity by positioning the apical domain, upstream of membrane-based polarity cues. These findings raised the question how vesicles acquire apicobasal directionality independent of polarized target membrane domains. Here, we show that the apical directionality of vesicle trajectories depends on intracellular actin dynamics during the establishment of membrane polarity in the *C. elegans* intestine. We find that actin, powered by branched-chain actin dynamics, determines the position of apical membrane components, PARs, and itself on expanding membranes. Using photomodulation, we demonstrate that F-actin travels through the cytoplasm and along the cortex towards the future apical domain. Our findings suggest an alternative polarity model where actin-dependent directional trafficking inserts the nascent apical domain into the growing membrane to partition its apicobasal domains.

## Main Text

Prevailing epithelial polarity models place extracellular- and membrane-/junction-based core polarity cues (e.g., the partitioning-defective PARs) upstream of cell-intrinsic events during the establishment of cell and membrane polarity^1^. Once these core polarity cues have determined the positions and identities of apicobasal membrane domains, these domains serve as target domains to recruit polarized membranous and submembranous molecules from inside the cell that expand and maintain them. For instance, the apicobasal directionality of intracellular biosynthetic trafficking that contributes the bulk of polarized membrane components is thought to strictly depend on cognate recognition sites at these polarized target domains^2^. Domain exclusion of the membrane-based core polarity cues (e.g., the PARs), initiated by cell-cell contact, is also considered a principle mechanism of modulating membrane polarity within the tissue context ^3^. Thus, whether during the establishment or modulation of membrane polarity, the membrane-based apicobasal core polarity cues are thought to impart the initiating polarizing event at the membrane. In these models, the process of polarized membrane biogenesis is viewed as an exchange of molecular components within membrane domains demarcated by membrane-based core polarity cues.

We previously identified several molecules in forward *C. elegans* tubulogenesis screens whose perturbation reversibly changed the position of the apical domain (lumen) in already polarized cells of the developing intestine, overriding the polarity established by the core polarity cues (apicobasal membrane polarity conversion). Some of the identified molecules were components of vesicular trafficking (membrane glycosphingolipids, clathrin/AP-1, SEC-23, and other components of the biosynthetic-secretory pathway ^4–7^. The analysis of this polarity phenotype revealed that vesicle-based polarity cues were required to specify the position of the apical domain on the expanding membrane, upstream of PARs, from the time of membrane polarity establishment in still dividing and migrating cells throughout further net polarized membrane addition required to maintain this polarity in postmitotic cells within the fixed but still growing tissue context. Based on these findings, we proposed an alternative mode of polarized membrane biogenesis where apical and basolateral membrane domains are partitioned by the asymmetric insertion of the apical domain via biosynthetic trafficking rather than by the molecular exchange of membrane components within domains previously demarcated by membrane-based core polarity cues ^7^. This mode of membrane polarization would also explain recent observations on related polarity phenotypes in different tissues and species (including humans) that also suggested a role of intracellular vesicular trafficking (albeit polarized recycling) upstream of membrane-based core polarity cues such as the PARs in the positioning and maintenance of apical/anterior membrane/cortex domains ^1, 8–13^.

However, it remained unclear how vesicular trafficking could be polarized without the prior polarization of target membrane domains, considered a prerequisite for polarized vesicles to reach their target. We suggested that a transient cytoskeletal structure might asymmetrically orient the biosynthetic-secretory pathway from the ER to the plasma membrane at the time of *de novo* polarized membrane biogenesis, regulated by cell intrinsic or extrinsic signals and oriented by membrane-based landmarks other than already polarized membrane domains ^7^. However, suitable cytoskeletal tracks to confer such long-range apical directionality to vesicles have not yet been identified in any system. Moreover, in the *C. elegans* intestine, neither microtubules, thought to contribute to, but not to establish, its polarity, nor actin, considered entirely dispensable for it, appeared to provide candidate cytoskeletal material (the function of non-polarized intermediate filaments in the intestine is thought to be structural^14, 15^. We here focus on a group of cytoskeleton-associated molecules that we identified by the same apicobasal intestinal polarity conversion in the same tubulogenesis screens that identified the vesicle-based polarity cues - making them candidate components of such a proposed cytoskeletal structure.

### The branched-chain actin modulators (bcAMs) UNC-60, ARX-2 and CAP-1 specify the position of the apical domain and lumen in embryonic and larval *C. elegans* intestinal cells

In RNAi-based genome-scale tubulogenesis screen we identified several genes whose knockdowns induced apicobasal membrane polarity conversion in the developing *C. elegans* intestine. Polarity conversion was defined as the basolateral displacement of apical membrane components and polarity cues in any or all the 20 cells of the single-layered *C. elegans* intestine, with and without subsequent ectopic basolateral lumen formation (transformation of the basolateral membrane into a junction-bounded apical membrane with microvilli ^5^. This full membrane polarity change can be tracked by the apical identity marker ERM-1 whose recruitment to the membrane indicates successful apical polarization ^7^. Among the identified molecules, 3 were the key components of the branched-chain actin machinery (here referred to as branched-chain actin modulators/bcAMs; Fig.1): UNC-60/cofilin (actin-depolymerizing), ARX-2/ARP-2/3 (actin nucleating) and CAP-1 (actin capping). A scaled-intensity RNAi approach (employed to generate a range of mild to moderate defects without disrupting cellular morphogenesis; Methods) revealed that ERM-1: (1) was displaced to basolateral domains and ectopic lumens in cells of the late-embryonic/larval intestine that is fully formed but still expands (mildly affected *unc-60-, arx-1-, cap-1(RNAi)* animals that arrest as larvae; Fig.1d-n); and (2) failed to be polarized to the apical domain in cells of the early-embryonic intestine, at the time when its definitive polarity and lumen position are established (pan-membranous ERM-1 in strongly affected *unc-60-, arx-1-, cap-1(RNAi)* animals that arrest as early embryos, before or during intestinal intercalation; Fig.1o-v). Thus, each of these 3 bcAMs is required to specify the position of the apical domain on not yet polarized membranes of pre-mitotic cells and on already polarized but still expanding membranes of postmitotic cells (see Fig.3S for intestinal development and net apical membrane expansion in embryonic and larval intestinal cells). bcAM loss also fully aborted apical membrane (lumen) expansion inside the single-cell excretory canal (Fig.1w). Germline deletions and tissue-specific RNAi confirmed these phenotypes, demonstrated that these genes’ function cell autonomously in intestinal polarity, and suggested that maternal product was required for this function (Fig.1S). Imaging of polarized membrane markers and functional studies in larval intestines showed that the loss of each of the 3 bcAMs mispositions multiple apical, but not basolateral, membrane components, without disrupting the integrity of apical junctions that secure the positions of apical and basolateral membrane domains (Fig.1, S1). Transmission electron microscopy (TEM) confirmed the presence of junction-bounded ectopic basolateral lumens with microvilli in *unc-60(RNAi)* larvae (Fig.1x-z). These features copy the apicobasal polarity phenotype induced by the loss of several trafficking molecules that were previously identified as apical polarity cues in these same tubulogenesis screens (e.g., glycosphingolipids/GSLs and CHC-1/clathrin; ^5, 6^, Introduction; Fig.S1 and below). We conclude that UNC-60, ARX-2, and CAP-1, like the previously identified apical polarity cues, are required to specify the position of the apical domain in the *C. elegans* intestine. The close similarity of the bcAM- and trafficking-dependent polarity phenotype made actin a candidate component of the proposed cytoskeletal structure that might direct vesicle trajectories to the membrane to asymmetrically insert the nascent apical domain (see Introduction, ^7^.

**Figure 1.**
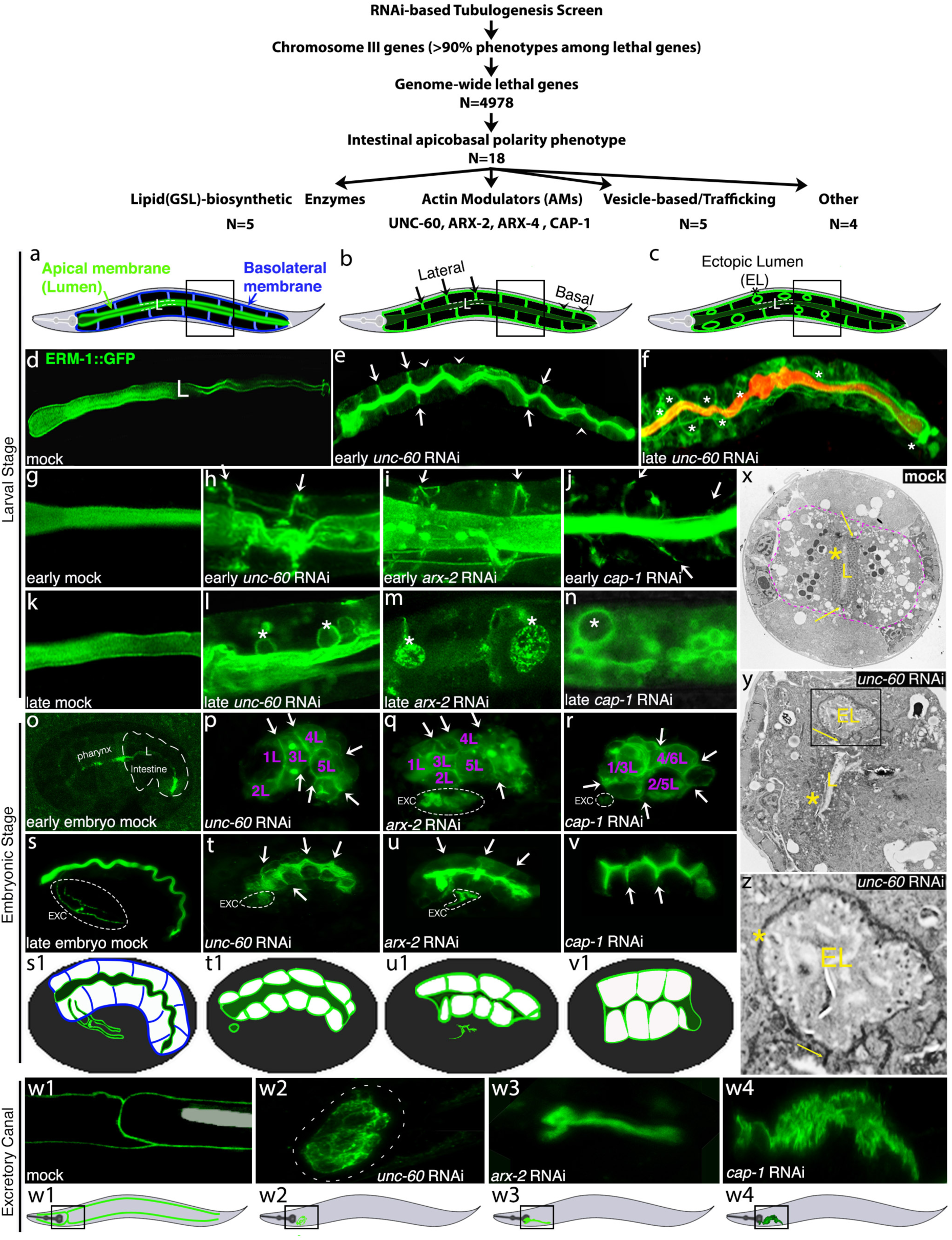
The *unc-60-, arx-2-, cap-1*-dependent apicobasal polarity phenotype. Genome-scale RNAi-based tubulogenesis screens identify 18 molecules whose reduction of function causes apicobasal polarity conversion in the developing *C. elegans* intestine. (a-c) Schematic of apicobasal polarity conversion on expanding membranes of larval intestines: (a) wild-type polarity, apical membrane: green, basolateral (BL) membrane: blue, (b) early stage polarity conversion with BL mislocalization of the apical domain, (c) late stage polarity conversion with formation of ectopic lumens (ELs). Note that ELs develop from BL membranes. Here and below, the lateral portion of the BL membrane is indicated by arrows, the basal portion by arrowheads. See Fig.S3 for *C. elegans* intestinal development. (d) Wild-type (mock) ERM-1::GFP identifies the apical domain (lumen). (e) BL ERM-1 mislocalization at early stage of polarity conversion (early), (f) formation of basolateral ELs (asterisks) at later stage of polarity conversion in *unc-60(RNAi)* larval intestines (late). Note that intralumenal DsRed bacteria (f) fill apical membrane blebs, but not basolateral ELs (nor ERM-1+ apical vacuoles; see text). (g-n) Magnified view of 1 or 2 pairs of opposing cells (corresponding to boxed areas in (a-c)) in *unc-60-, arx-2- and cap-1(RNAi)* larval intestines (g, k: wild-type). Note speckled ERM-1::GFP in ELs (m; asterisks) suggesting dysmorphic microvilli (MV; compare x-z). (o-r) Failure of ERM-1 polarization and intercalation arrest in early-embryonic *unc-60-, arx-2- and cap-1(RNAi)* intestines, (s-v) BL ERM-1::GFP mislocalization (arrows) in post-intercalation late-embryonic *unc-60-, arx-2- and cap-1(RNAi)* intestines (o, s: wild-type; identities of E progenitors in purple; see Fig.S3 for schematic of early-embryonic intestinal intercalation). (s1-v1) Corresponding schematics (note absence of excretory canal [EXC] lumen extension in all; compare w1-4). (w1-w4) Failure of intracellular apical membrane (lumen) expansion in *unc-60-, arx-2- and cap- 1(RNAi)* excretory canals. Corresponding schematics of whole worms are shown beneath (boxes indicate areas magnified above). (x-y) Transmission electron microscopic (TEM) image of wild-type (x) and *unc-60(RNAi)* (y and z) larval intestines (cross section of 2 cells in x and y, outlined by purple dashes in x): note main lumen (L) with dense MVs (asterisk) in x versus main lumen with disorganized MVs in y, and EL (boxed) above main lumen at basolateral membrane in y, magnified below. (z) EL has sparse dysmorphic MVs at apical (lumenal) membrane (asterisk). Yellow arrows point to intact apical junctions. Note that EL is separated from the main lumen by such a junction, distinguishing it as *bona fide* ectopic basolateral lumen.* Confocal images are shown throughout. *Distinction of apical domain phenotypes: Ectopic basolateral lumen: ERM-1+ apical membrane with microvilli and surrounding junctions, located within BL membrane domain; ectopic intracellular lumen: cytoplasmic ERM-1+ apical membrane-bound inclusion, with microvilli, no junctions; apical vacuole: cytoplasmic ERM-1+ apical membrane-bound inclusion, no microvilli, no junctions; apical membrane blebs: ERM-1+ membrane blisters contiguous with main lumen.

### Actin itself partitions apicobasal membrane domains and UNC-60, ARX-2 and CAP-1 interact with each other and actin in this function

Currently, actin is considered dispensable for *C. elegans* intestinal polarity ^15^ and ACT-5 is considered to be the sole intestinal actin ^16^ among 5 closely related *C. elegans* actins, encoded by 5 different genes: *act-1-5*; designated actin isoforms below for simplicity). Since actin is expected to operate downstream of bcAMs, we re-examined its role in polarity. The assumption of actin’s dispensability for polarity chiefly rests on chemical interference, balanced *act-5* germline deletions (containing maternal product) and on ACT-5-3’UTR RNAi (mild depletion) that result in late (L1-larval) lethality and retain the ability to polarize junctions ^14, 16^. We found that the widely used *act-5* double stranded RNA (produced by a clone of the Ahringer bacterial RNAi feeding library; Fig.2Sb) not only induced larval, but also early embryonic intestinal polarity defects and lethality (Fig.2a-d; note, however, that this RNA targets all 5 actin isoforms – we thus refer to it as *actin* RNAi below). In addition to the previously reported *act-5* dependent structural apical domain (lumen) morphogenesis defects in larval intestines (complete loss of microvilli ^16, 17^, *actin* RNAi also generated apical vacuoles and ectopic basolateral lumens indicative of polarity conversion (Fig.2e-g; see legend for distinction of: apical membrane blebs, apical vacuoles, intracellular ectopic lumens, and basolateral ectopic lumens). Mild *actin* RNAi (Methods) also mispositioned apical, but not basolateral, membrane components on expanding membranes of larval intestinal cells, without affecting junction integrity (Fig.1w). Moreover, *actin* RNAi induced in larvae (after polarity establishment) was sufficient to change the polarity of these already polarized membranes (basolateral-to-apical; Fig.2j), demonstrating that actin exerts a direct effect on *de novo* polarized membrane biogenesis (see Methods for the postmitotic larval intestine as a model for the *in vivo* analysis of *de novo* polarized membrane biogenesis). Thus, actin is required for intestinal membrane polarity and its loss copies the bcAM-induced polarity phenotype, consistent with actin’s role as downstream effector of bcAMs in polarity.

**Figure 2.**
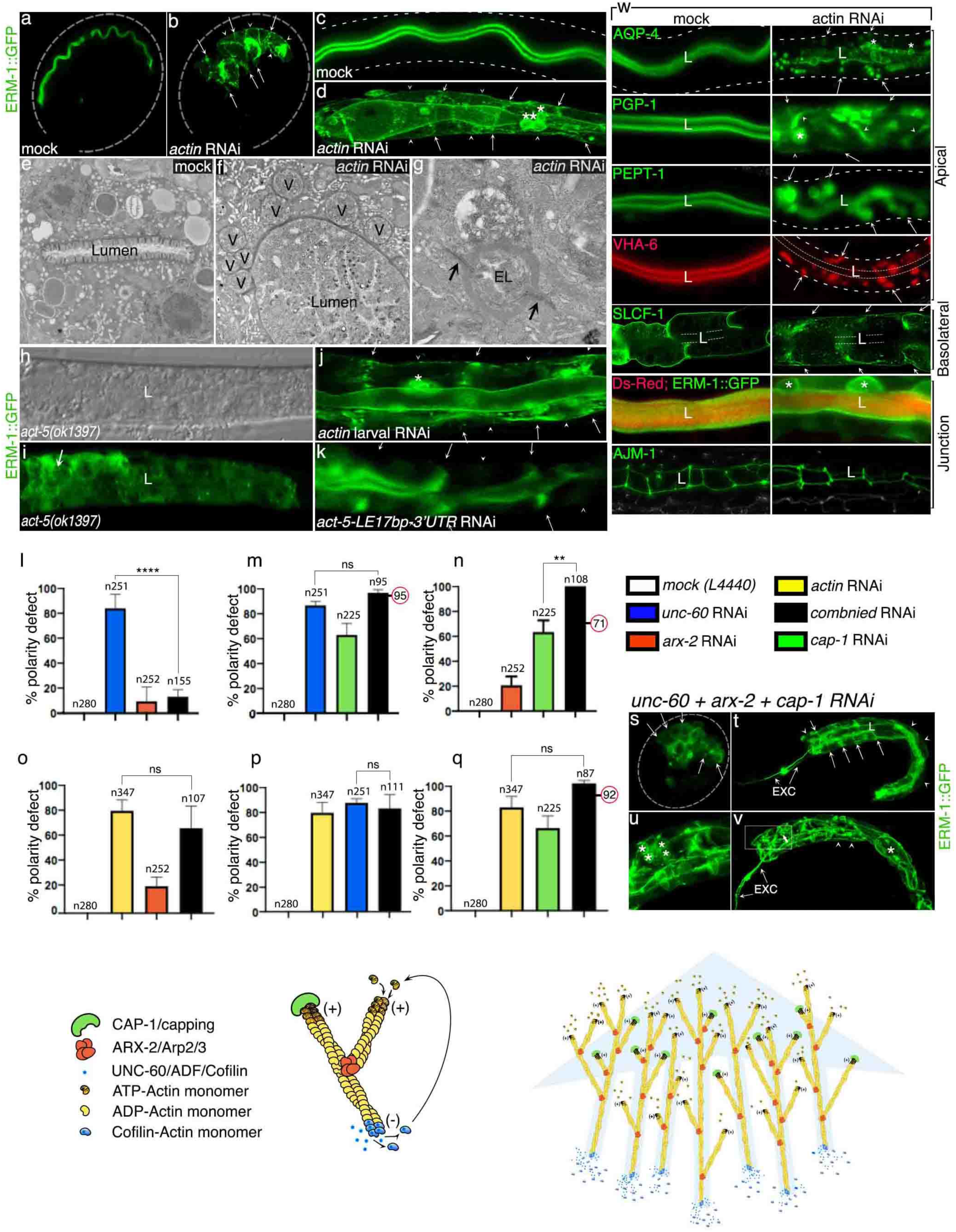
Actin partitions apicobasal membrane domains in the *C. elegans* intestine and functionally interacts with the 3 branched-chain actin modulators in polarity. (a) ERM-1::GFP is apical in mock RNAi embryonic intestines (eggshell dotted; see Fig.1 for schematic of embryonic intestine). (b) *act-5* RNAi (Ahringer dsRNA clone, designated *actin* RNAi since it targets all five actins; Fig.S2b) causes basolateral displacement of ERM-1::GFP. (c, d) Wild-type versus altered membrane polarity in *mock(RNAi)* versus *actin(RNAi)* larval intestines. Arrows in a-d indicate lateral displacement of ERM-1::GFP and arrowheads basal displacements (basal membrane indicated by dotted line in [c] since not visible in mock). Note lumenogenesis defects, cytoplasmic ERM-1::GFP displacement, ERM-1+ vacuolar inclusions and ectopic lumens (EL, asterisks) in (d). (e) TEM cross section of *mock(RNAi)* intestine shows intestinal lumen with intact microvilli, (f) dilated lumen with complete loss of microvilli in *actin(RNAi)* intestine (V(s): apical vacuoles), (g) magnified panel depicting basolateral ectopic lumen (EL) with surrounding intact junctions (arrows; note that actin is required for microvilli assembly, disqualifying microvilli as defining characteristic for ELs in this specific case, compare Fig.1 legend*). (h and i) Differential interference contrast (DIC)/Nomarski and confocal images of *act-5(ok1397)* (Fig.S2 for genotype): note full cytoplasmic ERM-1::GFP displacement from the lumen. (j) Conditional larval *actin* RNAi also induces basal and lateral displacement of ERM-1::GFP (arrowheads and arrows, respectively), as does *act-5-LE17bp-3’UTR* RNAi (k; see text and Fig.S2 for dsRNA). (l-q) Genetic interactions of actin and bcAMs during polarized membrane expansion. The *actin(RNAi)* polarity defect (yellow; o-q) is not significantly enhanced by the loss of bcAMs. >3 replicas were analyzed. Numbers in red circles (e.g., 95 or 71 in “m” and “n”) indicate additive values (=no interaction; for simplicity, statistical comparison [brackets] is shown w/o this adjustment; Methods). *t-test* for significance: **p= *p < 0.01* (*n*: total number of animals scored). (s-v) Triple *unc-60, arx-2, cap-1* RNAi dramatically increases the polarity defect: (s) early embryo. (t-v) L1-larva. Arrows: lateral, arrowheads: basal, ERM-1 displacement, asterisks: ectopic lumens/ELs. EXC: excretory canal. w panels: *actin* RNAi changes the positioning of apical, but not basolateral membrane components and maintains junction integrity (as does RNAi with bcAMs; Fig.S1, see there for details). L: lumen, dashed lines: intestinal basal border. Where not indicated otherwise, confocal images of sections of, or full (t,v), larval intestines are shown; 2 pairs of opposing cells in w. **Schematics.** Branched chain actin dynamics. Left: Capping, branch-nucleating, and disassembly activities of the 3 bcAMs at the actin filament. Right: Actin treadmilling. Repeat cycles of filament assembly and disassembly that generates the impression of filament movement (directionality indicated by blue shading).

Consistent with actin’s function in intestinal polarity and suggesting it was mediated by ACT-5, RNAi that predominantly targets the ACT-5 3’UTR (inclusive of 17 base pairs of exonic sequence, some also present in other actins; designated *act-5-LE17bp-3’UTR*; Fig.S2b) copied the *actin(RNAi)* polarity phenotype (Fig. 2k). In addition, in an *act-5* deletion mutant (a large deletion, including the deletion of 500bp promotor sequence) in a balanced background (i.e., in the presence of maternal product), ERM-1 was fully displaced from the lumen, supporting ACT-5’s role in apical domain biogenesis but masking possible effects on apical domain positioning (Fig.2h-i). Both the germline deletion and RNAi only induced larval, not embryonic lethality. We concluded that ACT-5 requires maternal product and/or additional actin isoforms for its presumed function in polarity.

To determine the relation of the 3 bcAMs to each other and to actin in this polarity function, we assessed genetic interactions among bcAMs and between bcAMs and actin. Experiments were performed by mild-to-moderate RNAi to avoid disrupting full gene function expected to mask specific effects on membrane polarity (see Methods for discussion of choice of experimental approach). Triple RNAi with all bcAMs enhanced the intestinal polarity phenotype of each (Fig.2s-v; enhancement of: early-embryonic intercalation-, lumenogenesis-, and membrane-polarity defects; and of larval polarity conversion and ectopic lumen formation). To assess bcAM/actin interactions during the process of polarized membrane biogenesis, we examined their ability to induce polarity conversion in postmitotic larval cells (see above for experimental model). Consistent with their opposing functions in actin filament assembly (Fig.2 schematic), the *unc-60* (filament depolymerizing) dependent polarity conversion was suppressed by RNAi with *arx-2* (branch nucleating), while the *arx-2* dependent polarity conversion was enhanced by RNAi with *cap-1* (filament capping). However, interference with each bcAM failed to significantly enhance the *actin(RNAi)* induced polarity conversion, consistent with actin’s function as downstream effector of bcAMs (note that this analysis only reflects the interaction of bcAMs/actin in *de novo* polarized membrane biogenesis; it can be separated from their interaction in lethality; Fig.22l-q and S2a).

We conclude that actin is required for polarity in the *C. elegans* intestine and cooperates with bcAMs in specifying the position of its apical domain (lumen). These findings could support the hypothesis that actin provides the cytoskeletal structure that might direct biosynthetic trafficking to the nascent apical domain during membrane polarization.

### UNC-60, ARX-2, CAP-1, and actin specify the polarized distribution of apical PARs on expanding membranes of pre- and postmitotic intestinal cells

Actin and bcAMs have conserved functions in apical cortex and junction modeling which includes apical domain assembly in the *C. elegans* intestine ^18, 19^. They also maintain apical/anterior PARs in epithelia and in the *C. elegans* zygote ^8^. bcAMs/actin might therefore affect polarity by these canonical functions that maintain the apical domain rather than by a proposed novel function in directing biosynthetic trafficking to position this domain. To distinguish between these possibilities and to assess the position of bcAMs/actin in the process of membrane polarization, we next examined their polarity function in relation to that of the PARs (bcAMs’ canonical function should operate downstream, their proposed novel function upstream, of apical PAR positioning and function). We assessed the polarized distribution of endogenously tagged PAR-3, PAR-6, and PKC-3, the 3 apical PAR complex components, with and without ERM-1::GFP, on bcAM/actin depleted intestinal membranes during embryonic and larval membrane expansion and polarization (Fig.3; Fig.S3 for intestinal development and net apical membrane addition during this time). PAR-3 is considered the earliest polarity cue in the establishment of polarity in most epithelia, including the *C. elegans* intestine, where it is also the only PAR shown to be strictly required for polarity ^20, 21^. Mild RNAi with *unc-60*, *arx-2*, *cap-1*, and *actin* was sufficient to displace apical PAR-6 and PKC-3 to basolateral membranes during *de novo* polarized membrane biogenesis in expanding postmitotic larval cells (Fig.3; PAR-3 expression weans at this stage). Moreover, the initial polarized distribution of apical PARs, including PAR-3, was also dependent on each bcAM and actin during polarity establishment in the intercalating early-embryonic intestine (pan-membranous PAR-3, -6 and PKC-3 or no recruitment to the apical domain in more severely affected intestines; Fig.3 and legend for details). The effect of bcAMs and actin on PARs’ polarized distribution copied that of the previously identified vesicle-based apical polarity cues, here shown in *chc-1(RNAi)* early-embryonic, and *let-767(RNAi)* larval, intestines (Fig.3; *chc-1* encodes the clathrin heavy chain, *let-767* a glycosphingolipid biosynthetic enzyme ^5, 6^. We conclude that bcAMs and actin are required to specify the position of apical PARs on not yet polarized membranes of still dividing and migrating intestinal cells and on already polarized but still expanding membranes of postmitotic cells within the fixed epithelium. These findings further support the proposed non-canonical function of bcAMs and actin in membrane polarity via directional trafficking that likewise operates upstream of PARs in intestinal polarity ^7^. Our findings also reveal that branched chain actin dynamics functions upstream of the to date earliest polarity cue in the establishment of epithelial polarity in the C*. elegans* intestine.

**Figure 3.**
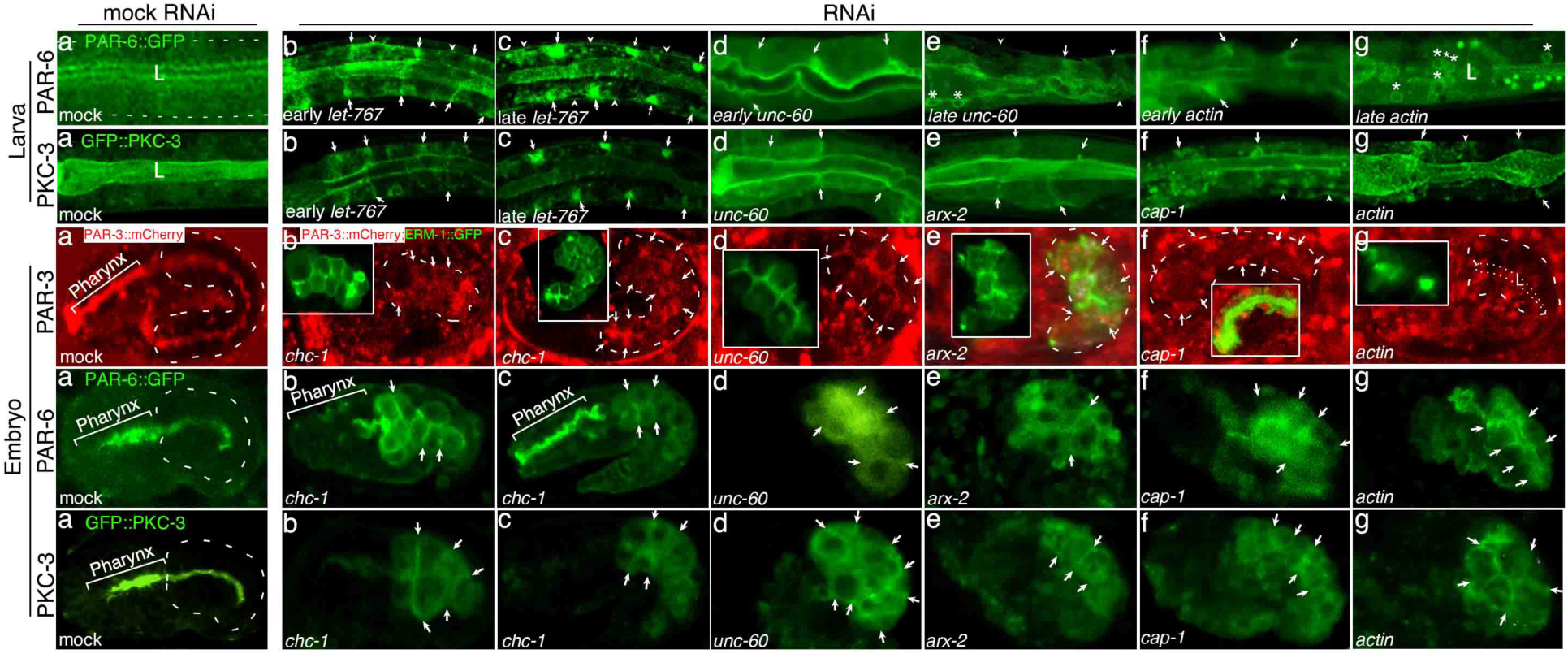
Branched chain actin dynamics functions upstream of apical PARs in intestinal polarity and interference with bcAMs/actin copies the effect of interference with trafficking-based apical polarity cues (*let-767* and *chc-1*; see text). The upper 2 rows show portions of early larval intestines, the lower 3 rows full embryonic intestines. (**1^st^ row**, a) PAR-6::GFP is apical in wild-type larval intestines (mock RNAi; no PAR-6 at basal membrane [indicated by dashed line]). (1^st^ row, b, c) lateral (arrows) and basal (arrowheads) PAR-6 displacement at early and later stages of the *let-767(RNAi)* polarity change (Fig.S3 for net apical membrane expansion in larvae). (1^st^ row, d, e) basolateral (arrows) and cytoplasmic PAR-6 displacement at early and late stage of *unc-60(RNAi)* polarity conversion; note ectopic lumen/EL development [asterisks]). (1^st^ row, f, g) *actin* RNAi induces basolateral (arrows) and cytoplasmic PAR-6 displacement (early and late stages; note PAR-6 displacement to apical and basolateral membrane blebs and ELs in (g) [asterisks], see text). (**2^nd^ row**, a) GFP::PKC-3 is apical in wild-type larval intestines (mock RNAi; no PKC-3 at basal membrane). (2^nd^ row, b, c) PKC-3 displacement during *let-767(RNAi)* polarity conversion. (2^nd^ row, d-g) RNAi with each of the 3 bcAMs and actin cause basolateral (arrows) and basal (arrowhead) PKC-3 displacement. (**3^rd^ row**, a) PAR-3::mCherry is apical in wild-type embryonic intestines (mock RNAi; no PAR-3 at the basal membrane [indicated with dashed line]; note PAR-3’s additional location at apical junctions). Intestines are double labeled with PAR-3::mCherry and ERM-1::GFP, the latter is also strictly apical in wild-type. PAR-3 assembles at the future apical domain in the pre-intercalation early embryonic intestine^20^; late wt embryo shown here) (3^rd^ row, b, c) PAR-3 is displaced to all sides of the *chc-1(RNAi)* early embryonic (pre-intercalation; bean stage) and late embryonic (post-intercalation) intestine (arrows point to basolateral membranes). Insets shows the corresponding pan-membranous mislocalization of apical ERM-1::GFP (sizes adjusted for clarity). (3^rd^ row, d-f) RNAi with each of the 3 bcAMs causes basolateral (arrows) PAR-3 and ERM-1 displacement (for clarity, PAR-3/ERM-1 overlap [light green] shown for *arx-2* RNAi and in the inset for *cap-1* RNAi). (3^rd^ row, g) RNAi with actin prevents PAR-3 recruitment to the apical domain (indicated by dotted line and L/lumen). Pre-intercalation intestines shown in d, e, g, post-intercalation in f. (**4^th^ and 5^th^ rows**, a) PAR-6::GFP and GFP::PKC-3 are apical in wild-type embryonic intestines (mock RNAi; no PAR-6 and PKC-3 at the basal membrane [indicated by dashed line]; note additional junction location of PAR-6/PKC-3). (b, c) PAR6/PKC-3 displacement in early embryonic (intercalating) and late embryonic (post-intercalation) *chc-1(RNAi)* intestines (note concomitant intercalation defects). (d-g) RNAi with each of the 3 bcAMs and actin prevents apical polarization of PAR-6 and PKC-3 (note intercalation and lumenogenesis defects; no lumen formation except in g [PAR-6]). Pre-intercalation intestines shown in all except g (PAR-6). Confocal images shown throughout. See Figs.1 and S3 for schematics of larval and embryonic intestines, intestinal intercalation, and net addition of apical membrane during intestinal development. All fluorophore fusion proteins are expressed from their germline loci (Tab S2).

### UNC-60, ARX-2, CAP-1, and actin concurrently shift their positions from the basolateral to the nascent apical domain during polarity establishment

Consistent with a putative function of bcAMs in guiding vesicles to the plasma membrane, bcAMs have been associated with anterograde vesicle trajectories in non-polarized cells (an association conversely interpreted as the vesicle-based delivery of bcAMs to the cortex ^22^. However, in the *C. elegans* intestine, the location of ACT-5, UNC-60, ARX-2, and CAP-1 is thought to be restricted to the apical domain (refs), an unlikely position from which to direct vesicles through the cytoplasm. We therefore first assessed whether bcAMs’ and actin’s location were compatible with their proposed non-canonical function and analyzed their intestinal expression pattern from the early embryo to adulthood. To resolve the subcellular localization of these ubiquitous molecules from the earliest stages of intestinal (E-cell) development (Fig.S3), we expressed UNC-60-, ARX-2- and CAP-1::GFP fusions under the pan-intestinal promotor *elt-2* (see Fig.4 legend for rationale of generating exogenously rather than endogenously tagged bcAMs). In cells of the pre-intercalating intestine, all 3 bcAMs were located in the cytoplasm and enriched along the nucleus and all sides of the membrane, UNC-60, and CAP-1 in a mesh-like structure (Fig.4). ARX-2 displayed a conspicuous speckled pattern throughout the cytoplasm and along membranes. Remarkably, during membrane polarity establishment, the location of all 3 bcAMs shifted from this pan-membranous to their definitive apical location, with ARX-2::GFP subsequently also labeling apical junctions (Fig.4). Consistent with the occupation of the same subcellular space by all 3 bcAMs, endogenously tagged ARX-2::RFP overlapped with CAP-1::GFP during its polarity shift and at the mature apical membrane. We conclude that bcAMs’ dynamic subcellular expression pattern is compatible with a developmentally regulated function of branched-chain actin dynamics in the regulation of membrane polarity that might include the apical routing of vesicles.

**Figure 4.**
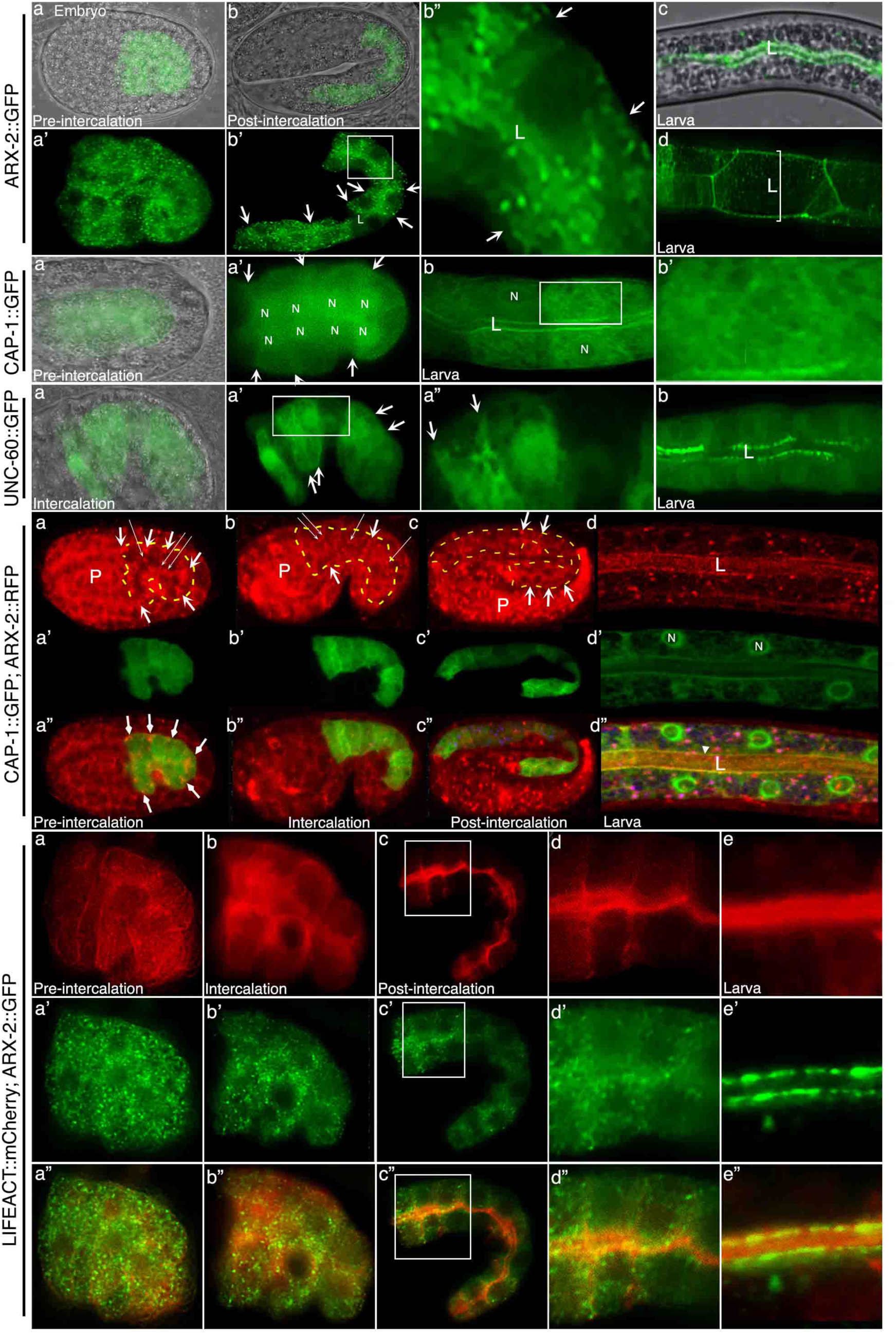
Dynamic subcellular localization pattern of UNC-60, ARX-2, CAP-1, and actin in the developing intestine. ARX-2::GFP: (a, a’) Pre-intercalation early-embryonic intestine (pre-bean stage): ARX-2 puncta throughout the cytoplasm and along all membranes; (b, b’) post-intercalation embryonic intestine (2-fold stage): ARX-2 puncta along all membranes (arrows point to lateral portions of basolateral membranes), enriched at the apical domain (lumen; L); (b”) magnified panel from b’; (c) expanding larval intestine: ARX-2 at the apical membrane (lumen; L); (d) mature adult intestine: ARX-2 at the apical membrane (lumen; L) and its peri-lumenal junctions (bracketed). **CAP-1::GFP**: (a, a’) Pre-intercalation early-embryonic intestine: CAP-1 is mainly cytoplasmic, with perinuclear (N=nucleus) and pan-membranous enrichment (arrows point to lateral membranes); (b) expanding larval intestine: CAP-1 remains cytoplasmic and becomes successively enriched at the apical membrane (lumen; L; no more basolateral membrane enrichment); a mesh-like structure is discernable, see higher magnification of inset in (b’). **UNC-60::GFP**: (a-b) UNC-60 also shifts from a cytoplasmic/basolateral (arrows) to a definitive apical location (lumen; L) during intestinal development; localization is similar to CAP-1 (above), with more pronounced mesh-like structure in the cytoplasm of intercalating cells (enlarged in a”). **ARX-2::RFP; CAP-1::GFP**: (a-d”) CAP-1 covers the area of ARX-2::RFP during intestinal development. Note hot spots of CAP-1 and ARX-2 overlap at the basolateral membrane angle (a-a”, arrows) in the early-embryonic intestine, and at the apical domain (arrowhead in d”) in the larval intestine; no colocalization of ARX-2 with CAP- at the nucleus (N) in larval intestines. Outline of intestine indicated by dashed line in b-d, basolateral membranes by arrows in b and c. Note that the *arx-2p-arx-2-rfp* germline insertion^53^ recapitulates the subcellular intestinal localization of transgenic *elt-2p-arx-2-gfp* (shown above), but puncta (long thin arrows in a and b) are not easily resolved due to low and ubiquitous endogenous *arx-2p-arx-2-rfp* expression (confocal image acquisition is substantially increased to identify ARX-2::RFP speckles, hence the increased background). **LifeAct::mCherry; ARX-2::GFP**: Actin is cytoplasmic and membrane-associated and copies the 3 bcAMs’ developmental shift from cytoplasmic/pan-membranous to apical. ARX-2 puncta decorate LifeAct-positive membranes from the pre-intercalation (location pan-membranous; a– a”) to the mature intestines (location apical; lumen [e–e”]). Images of thin confocal sections and of Nomarski/confocal overlays are shown. Gain increased for imaging of ARX-2::RFP to resolve subcellular pattern. See Methods for generation of fluorophore fusions.

To optimize live imaging of F-actin dynamics throughout intestinal development, capture any putative intestinal actin (beyond ACT-5), and avoid interference with function, we generated *elt-2*-directed LifeAct::GFP, an exogenous actin-binding molecule devoid of appreciable phenotypic effects (see Methods for choice of exogenous rather than endogenous actin binding molecules). LifeAct revealed that F-actin recapitulates bcAMs’ dynamic developmental subcellular expression pattern: it is also present in the cytoplasm and enriched at all sides of non-polarized membranes in early-embryonic intestinal cells, from which it shifts location to the apical domain (lumen) - its destination - during membrane polarity establishment (Fig.4). Double staining illuminated the concomitant basolateral-to-apical location change of actin and bcAMs, with ARX-2 speckles colocalizing with actin within the cytoplasm and alongside the cell cortex throughout its shift from pan-membranous to apical. We conclude that the 3 bcAMs and actin undergo a basolateral-to-apical positional shift during polarity establishment that could support their proposed function in polarity by physically directing vesicle trajectories towards the expanding apical domain.

### All 3 branched-chain actin modulators and actin interact with vesicle-based apical polarity cues in apical domain positioning

Prior data suggested that the directionality of the anterograde biosynthetic-secretory pathway, required to supply all sides of the membrane, becomes fixed from the ER to the nascent apical domain during *de novo* polarized membrane biogenesis in pre- and postmitotic *C. elegans* intestinal cells ^7^. A postulated cytoskeletal structure that confers such directional constraint to biosynthetic trafficking is expected to be short-lived and dynamic. The basolateral-to-apical bcAM/actin shift during polarity establishment would satisfy this requirement, and bcAMs’/actin’s role in apical domain positioning mirrors that of the previously identified apical vesicle cues (Fig.5a; see above). Three lines of evidence further supported the hypothesis that the bcAM/actin-dependent intestinal polarity phenotype depends on trafficking: (1) the previously noted reversibility of the *act-5(RNAi)* lumenogenesis defect and lethality^16^; less consistent with a structural morphogenesis defect but consistent with the reversible trafficking-dependent polarity defect and lethality ^5^; (2) the bcAM/actin-dependent membrane polarity conversion in the presence of intact junctions (as above); (3) cold suppression of the bcAM/actin-dependent membrane polarity conversion at temperatures expected to chiefly inhibit vesicle movement (Fig.5b-c).

**Figure 5.**
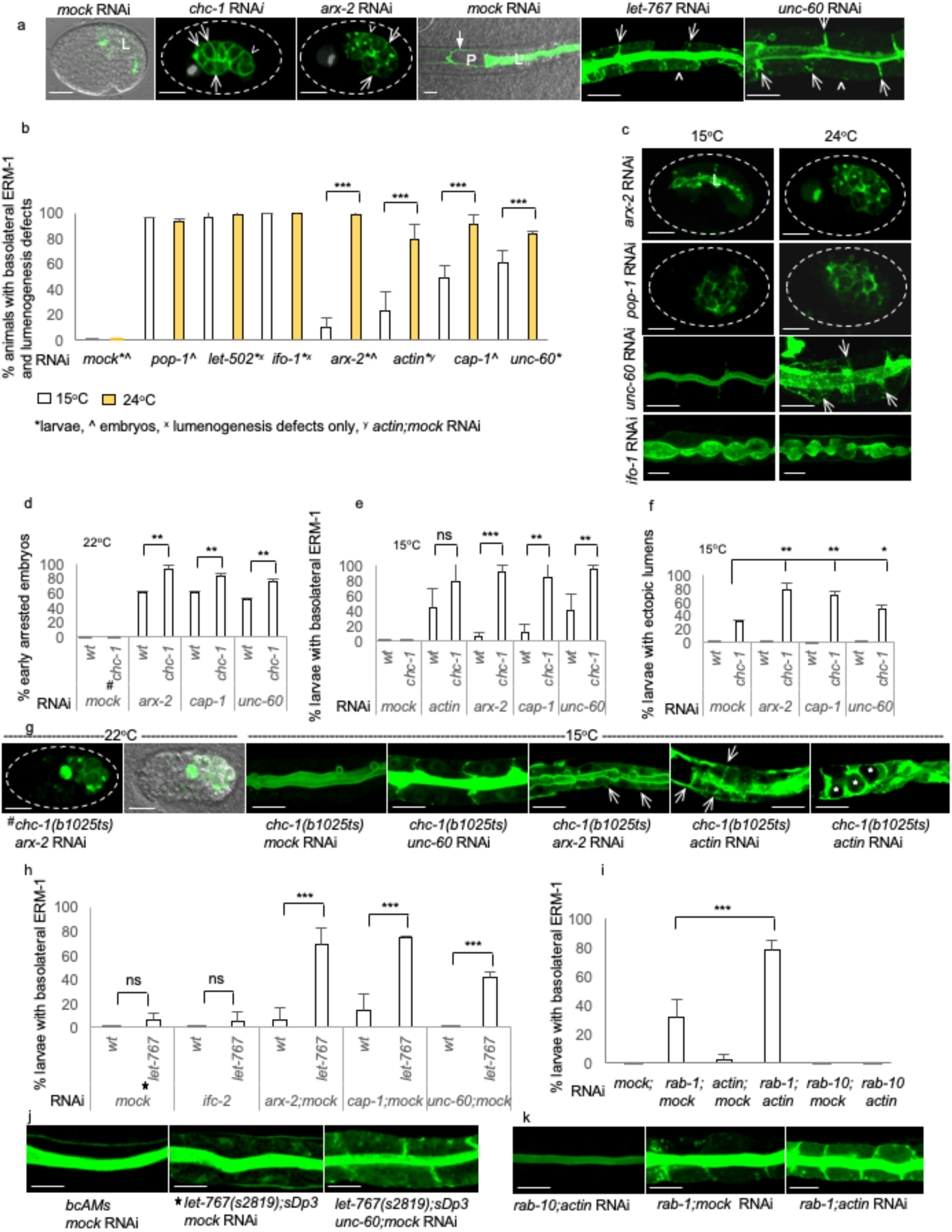
UNC-60, ARX-2, CAP-1, and actin interact with vesicle-based polarity cues in determining the position of the apical domain. (a) Failure of membrane polarization in *chc-1-* and *arx-2(RNAi)* embryonic intestines and basolateral-to-apical polarity conversion in *let-767* and *unc-60(RNAi)* L1-larval intestines. ERM- 1::GFP shown throughout. Here and below, arrows point to lateral, and arrowheads to basal portions of basolateral membranes. Embryo outline is dotted, excretory canal greyed out in embryos for clarity. L=lumen (apical domain). Scaled intensity RNAi approach was used here and below (Methods). (b-c) Cold suppression of the bcAM/actin-dependent polarity conversion. Basolateral ERM-1 displacement was assessed in larvae (*unc-60-, actin* RNAi), larvae and embryos (*arx-2* RNAi), or embryos (*cap-1, pop-1* RNAi), lumenogenesis defects in *let-502-* and *ifo-1(RNAi)* larval intestines. Note that similar polarity and lumenogenesis defects induced by interference with cell division/fate specification [*pop-1*/beta-catenin complex component], signaling [*let-502*/rho- dependent kinase] or with other structural cytoskeletal components [*ifo-1*/intermediate filament regulator] are not cold suppressed. Same stage animals were evaluated at different time points to account for delay in polarized membrane biogenesis at lower temperature. Suppression of polarity conversion is expected to occur via inhibition of the vesicle-based misrouting of apical membrane components. (c) Representative examples for embryonic and larval phenotypes. Note that at 15°C, lumen (L) is partially assembled in *arx-2(RNAi)* embryos and *unc-60(RNAi)* larval polarity conversion is almost completely suppressed. (d-f) Mild depletion of all 3 bcAMs and actin enhance the *chc-1(b1025ts)/*clathrin dependent lethality and polarity phenotypes at the permissive and restrictive temperature (see text). (g) Representative examples of double mutant/RNAi phenotypes. bcAMs/actin are titrated to levels that induce polarity defects in only few or no animals (no image shown). Large ERM-1::GFP+ vacuole in *chc-1(b1025)/arx-2* mutant/RNAi embryo (left side) corresponds to cystic excretory canal. *chc-1(b1025ts)/actin* mutant/RNAi larval intestines (right side) display a full synthetic ectopic lumen phenotype (*actin* RNAi is titrated to levels that do not induce any polarity defect on their own and *chc-1(b1025)* intestines display no full ectopic lumens at 15°C). (h-k) Mild depletion of each of the 3 bcAMs and actin induce synthetic polarity conversion in *let-767(s2819); sDp3* wild-type appearing larval intestines and enhance the *rab-1(RNAi)* larval polarity conversion (see text; corresponding representative images shown in j, k). *ifc-2* and *rab-10* are included as unrelated cytoskeletal and trafficking genes, respectively. Confocal and Nomarski/confocal overlay images of full embryos and 2 pairs of opposing larval intestinal cells are shown throughout. All values are mean ± SD, n >/= 3, p values calculated by two-tailed t test: p * < 0.05, **< 0.001, ***< 0.0001.

We next examined genetic interactions between these vesicle- and cytoskeleton-based polarity cues in polarity. Since null mutants cannot be assessed (see Fig.S2 legend for additional reasons to avoid full loss-of-function conditions), we titrated RNAi conditions and used temperature-sensitive and balanced mutants to test the expected ability of double mutant/RNAi to generate enhanced or synthetic polarity defects in a background of mild or no phenotypes, respectively.

Clathrin, a post-Golgi vesicle coat component and apical polarity cue (ref), was examined in the temperature-sensitive mutant *chc-1(b1025ts)* that exhibits early embryonic lethality (pre intestinal polarization) at temperatures above 22^0^C, but is viable with minimal basolateral ERM-1 displacement and ectopic lumens at the permissive temperature (15^0^C; Fig.5g). We titrated RNAi with bcAMs/actin, whose polarity defects are already suppressed in the cold (see above), to a level resulting in early embryonic lethality in only ½ or less than ½ of animals, and in no or almost no polarity defects during larval membrane expansion at 15^0^C (Fig.5d-f). These conditions enhanced the *chc-1(b1025ts)* early embryonic lethality and late ectopic lumen formation at the restrictive and permissive temperature, respectively, and synthetically induced basolateral ERM-1 displacement in all double *chc-1(b1025ts)*/*bcAM/actin(RNAi*) larval intestines at 15^0^C.

Glycosphingolipids (GSLs), Golgi and post-Golgi endomembrane-based apical polarity cues (ref), were examined in the *let-767(s2819)* mutant, balanced with the free duplication *sDp3*, that exhibits a full polarity/ectopic lumen phenotype in the absence of the duplication, but appears wild-type in its presence (Fig.5j; *let-767* is a GSL biosynthetic enzyme ^5^. All double GSL/bcAM/actin loss-of-function combinations either enhanced or synthetically induced polarity conversion in larval intestines (Fig.5h,j). Finally, *actin* RNAi, titrated to induce no polarity defect on its own, enhanced the basolateral displacement of ERM-1 on expanding larval intestinal membranes mildly depleted of RAB-1, a key component of the early secretory pathway that also functions as apical polarity cue (Fig.5i,k; ^7^.

We conclude that all components of the branched-chain actin machinery and actin functionally interact with the previously identified pre-Golgi-, Golgi- and post-Golgi-based apical polarity cues in apical domain positioning, consistent with the proposed non-canonical, trafficking-based function of actin dynamics in membrane polarity.

### Branched-chain actin dynamics confers apical directionality to pre- and post-Golgi vesicles during membrane polarization

If bcAMs/actin were to specify cellular membrane polarity by asymmetrically inserting newly-synthesized apical membrane components into the expanding membrane (proposed model; Fig.6a), the apical directionality of vesicles should depend on bcAMs/actin at the time of *de novo* polarized membrane biogenesis. Indeed, vesicular (and nuclear) repositioning has been noted to occur during the establishment of *C. elegans* intestinal polarity and its contribution to polarity establishment has been hypothesized but not yet demonstrated ^23, 24^. The visual tracking of biosynthetic vesicles from the ER to the plasma membrane in single cells has been difficult in any system, since biosynthetic cargo is thought to be successively enriched on its way to the plasma membrane while vesicle-coat and vesicle-membrane components are consecutively shed and recruited on this route ^25, 26^. To start mapping vesicle trajectories through *C. elegans* intestinal development, we directed expression of the ubiquitous RAB-11 (marking endosomes with presumed apical directionality) and RAB-10 (marking endosomes with presumed basolateral directionality to the intestine by *elt-2* to visually resolve their subcellular positions in both pre- and post-intercalation intestinal cells (^27^; Methods). We found that RAB-11+ post-Golgi vesicles, previously identified as apical polarity cues ^7^, indeed exhibit a bcAM/actin synchronous positional shift from the cytoplasm to the apical domain in the developing wild-type intestine, consistent with a directional change of presumed apical vesicle trajectories during membrane polarity establishment (Fig.6b). Based on this finding we next examined whether the positioning of these and other, not yet well mapped, vesicle populations was dependent on bcAMs/actin during *de novo* polarized membrane biogenesis.

**Figure 6.**
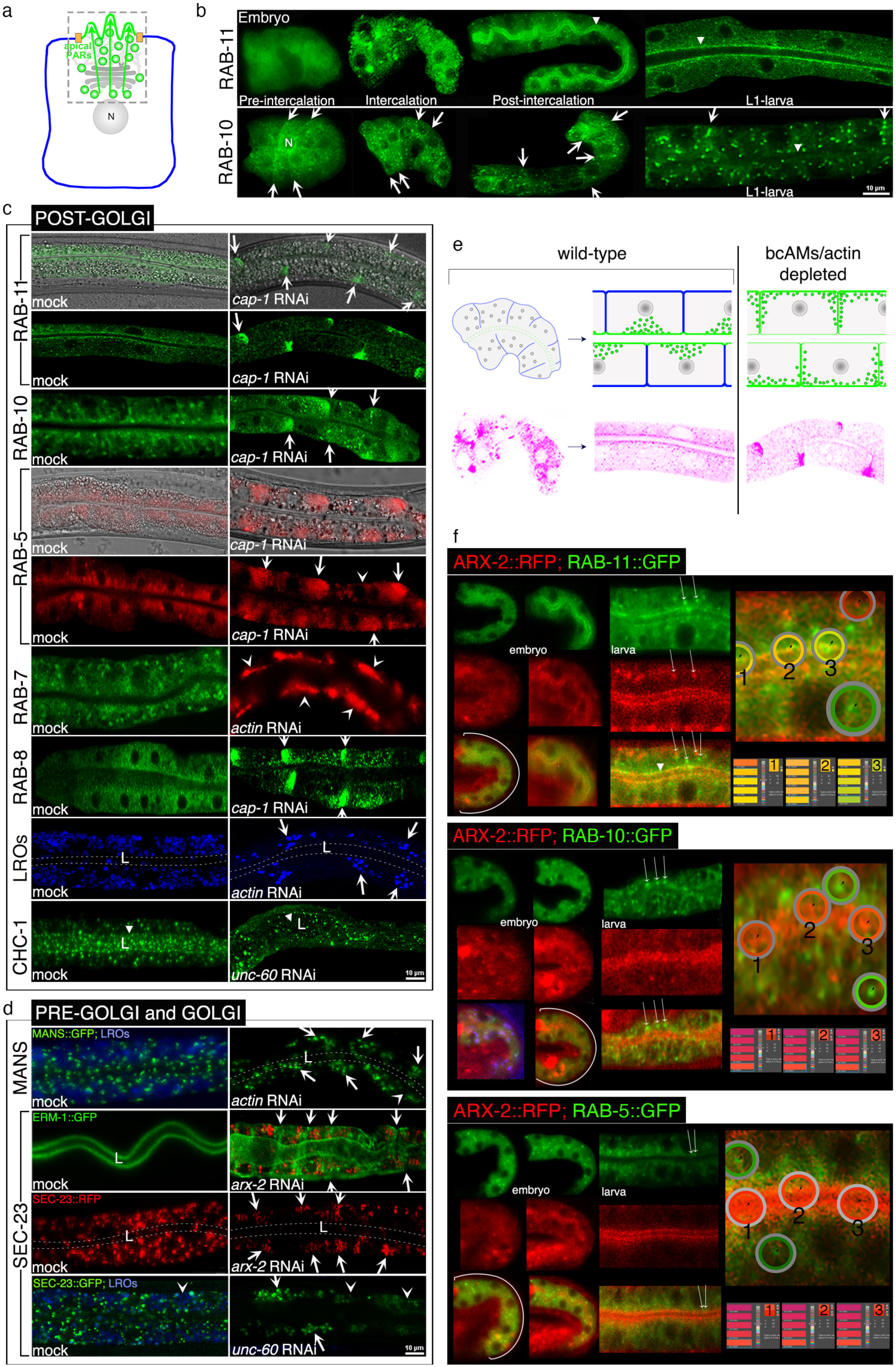
The apical targeting of pre- and post-Golgi vesicles during *de novo* polarized membrane biogenesis depends on UNC-60, ARX-2, CAP-1, and actin. (a) The proposed apicobasal polarity model (see text;^7^: the anterograde biosynthetic-secretory pathway is constrained towards the expanding apical domain and partitions apical and basolateral membrane domains; apical PARs and junction components are concomitantly or subsequently recruited to the apical domain, with apicolateral junction assembly securing the expanding apical domain (not shown; mode of PAR delivery is not determined). Side view of intestinal cell is shown, dashed box contains dynamic components during polarity establishment. Apical endo-, plasma membranes and PARs: green; basolateral membranes: blue; junctions: orange; Golgi and proposed cytoskeletal structure: grey, N=nucleus. (b): Developmental subcellular intestinal expression profile of RAB-11::GFP and RAB-10::GFP. Recruitment of RAB-11 and RAB-10 to vesicle membranes before and during intercalation, followed by strong apical (RAB-11) and mild pan-membranous (RAB-10) enrichment. (c-d) Apical components of post-Golgi-, pre-Golgi- and Golgi endomembranes/vesicle populations fail to reach the lumen (L, arrowhead, indicated by dashed lines where difficult to distinguish from background) and are mislocalized to basolateral (arrows) and basal (arrowheads) membranes of expanding *unc-60-, arx-2-, cap-1 and actin(RNAi)* late-embryonic/early-larval intestinal cells (see Fig.S4 for additional results and controls). RAB-11, CHC-1/clathrin and Sec-23/COPII are vesicle-based apical polarity cues (see text; ^7^). Corresponding Nomarski images (shown for RAB-11 and RAB-5) demonstrate morphologically intact intestines. ERM-1::GFP displacement (shown for SEC-23::RFP) reveals concomitant change in membrane polarity. (e) Schematic of polarized distribution of vesicles with apical destination during polarity establishment in expanding wild-type versus bcAM/actin depleted intestinal cells. Wild-type pre-intercalation intestine shown to the left, magnified section of 2 pairs of opposing wild-type versus bcAM/actin depleted cells of post-intercalation intestine to the right. Pink vesicles beneath trace RAB-11 vesicles from b and c. Schematic is simplified and only shows enrichment of apical vesicles (green) along membranes. Note concomitant change in membrane polarity. (f) Colocalization experiments with ARX-2::RFP and RAB-11::GFP, GFP::RAB-10 and ::RAB-5 during intestinal polarization. Panels on the right show magnified merged portions of double-labeled intestines, measured by the Eyedropper tool (Adobe-22.4.3 DIC color guide), revealing lack of direct overlap (yellow) at vesicles (long thin arrows) for all combinations, except for apical RAB-11 at the lumen (full arrowhead; see Fig.S4 for additional results). Cytoplasmic overlap of all markers in embryonic intestines indicated by brackets. Confocal and confocal/Nomarski overlay images of full early-embryonic and partial late-embryonic and early-larval intestines (2 pairs of opposing cells) are shown. Background increased in ARX-2::RFP intestines to resolve ARX-2 speckles (compare Fig.4). All labeling via fluorescent fusion proteins (Methods). Blue distinguishes autofluorescent lysosome related organelles (LROs). See Methods for generation of fluorophore fusions.

We assessed the subcellular location of pre- and post-Golgi vesicle populations during net apical membrane addition in postmitotic larval intestinal cells (see Methods and Fig.3S for experimental model). We found that mild interference with any of the 3 bcAMs and actin, but not with *sma-1*/spectrin (another component of the apical cytoskeleton), dramatically perturbed the pan-cytoplasmic wild-type distribution of multiple vesicle populations (Figs.6, S4). Mild depletion of each bcAM or actin not only misdirected RAB-11+ vesicles to basolateral membrane domains, but resulted in the basal/basolateral mislocalization of RAB-10+ recycling-, RAB-5+ early-endocytic-, RAB-7+ late-endocytic-, and RAB-8+ secretory endosomes, as well as of lysosome-related-organelles (LROs), all of which failed to reach the apical domain in otherwise intact appearing intestines (Figs.6c, S4A; note that the trafficking defect is specific [apical-to-basolateral position change] rather than general [cytoplasmic retention]). *bcAM/actin* RNAi also prevented the apical enrichment of clathrin-coated (CHC-1+) post-Golgi vesicles, tubulated MANS+ Golgi membranes and directed COPII-coated (SEC-23+) pre-Golgi vesicles to basal domains (Figs.6d, S4B; pre-Golgi vesicles and Golgi mini stacks are also distributed throughout the cytoplasm of *C. elegans* intestinal cells ^27^). The vesicle coat components SEC-23 and CHC-1 were also previously identified as polarity cues in the *C. elegans* intestine ^6^. We conclude that UNC-60, ARX-2, CAP-1, and actin are required for the apical positioning of all tested pre- and post-Golgi vesicle populations during *de novo* polarized membrane biogenesis, including those harboring vesicle-based apical polarity cues. These findings are consistent with a model where branched-chain actin dynamics asymmetrically directs anterograde vesicle trajectories from the ER to the nascent apical domain during membrane polarization (Fig.6a,e). Our findings could also suggest that multiple and presumably highly specialized distinct vesicle populations are co-opted during *de novo* polarized membrane biogenesis to deliver apical membrane components.

We considered that points of overlap between the 3 bcAMs and vesicles should identify sites from which bcAMs might direct vesicles and thus reveal the underlying mechanism for this process. For instance, bcAMs might power actin comets or plumes generated at specific vesicles for forward propulsion, a possibility suggested by the speckled pattern of ARX-2 (Fig.4). A colocalization analysis of bcAMs and post-Golgi vesicles throughout development revealed their proximity in the pre-intercalation intestine but failed to detect any direct overlap during *de novo* membrane biogenesis in larval intestines, except for ARX-2::RFP colocalization with RAB-11+, and, occasionally, RAB-5+ and RAB8+ vesicles, at the apical domain (Figs.6f, S4C-D). These results suggested that bcAMs, rather than propelling specific vesicles via vesicle-based actin assemblies, might direct entire vesicle populations by powering broader actin structures, consistent with their ability to confer directionality to multiple pre- and post-Golgi vesicles.

### A transcytotic basolateral-to-apical actin shift during membrane polarity establishment in the *C. elegans* intestine

To visually identify such an actin structure, we examined cells of the developing *C. elegans* intestine for the presence of actin assemblies with documented roles in directional trafficking. Obvious actin cables, the classical actin structure for directional vesicle transport via atypical myosins, appeared to be missing. However, short, membrane-close F-actin trails, implicated in apical secretion in other epithelia ^28, 29^, might not have been resolved by high resolution confocal or electron microscopy ^17, 30^. To image the apical submembranous intestinal actin cytoskeleton at molecular resolution, we employed stochastic optical reconstruction microscopy (STORM) in thinly mounted L1-larvae using phalloidin (Fig.8A, legend, Methods). This approach was able to resolve single actin microfilaments in microvilli of expanding apical intestinal membranes at nanometer resolution (average 17 filaments/microvillus; Fig.8Aa). Double-color STORM of actin and ERM-1::GFP (the latter detected by anti-GFP nanobodies) revealed full overlap of ERM-1 with actin in microvilli (Fig.8Ac; confirming immuno-TEM studies ^17^. However, neither phalloidin+ nor ERM-1+ trails emanated from a peri-lumenal actin belt postulated to root microvillar actin. In fact, no such belt was identified, nor junctional actin. In contrast, double-color STORM of actin and IFB-2 (detected by anti-MH33 nanobodies) revealed the rooting of actin filaments in a peri-lumenal intermediate-filament lattice (Fig.8Ab).

**Figure 7.**
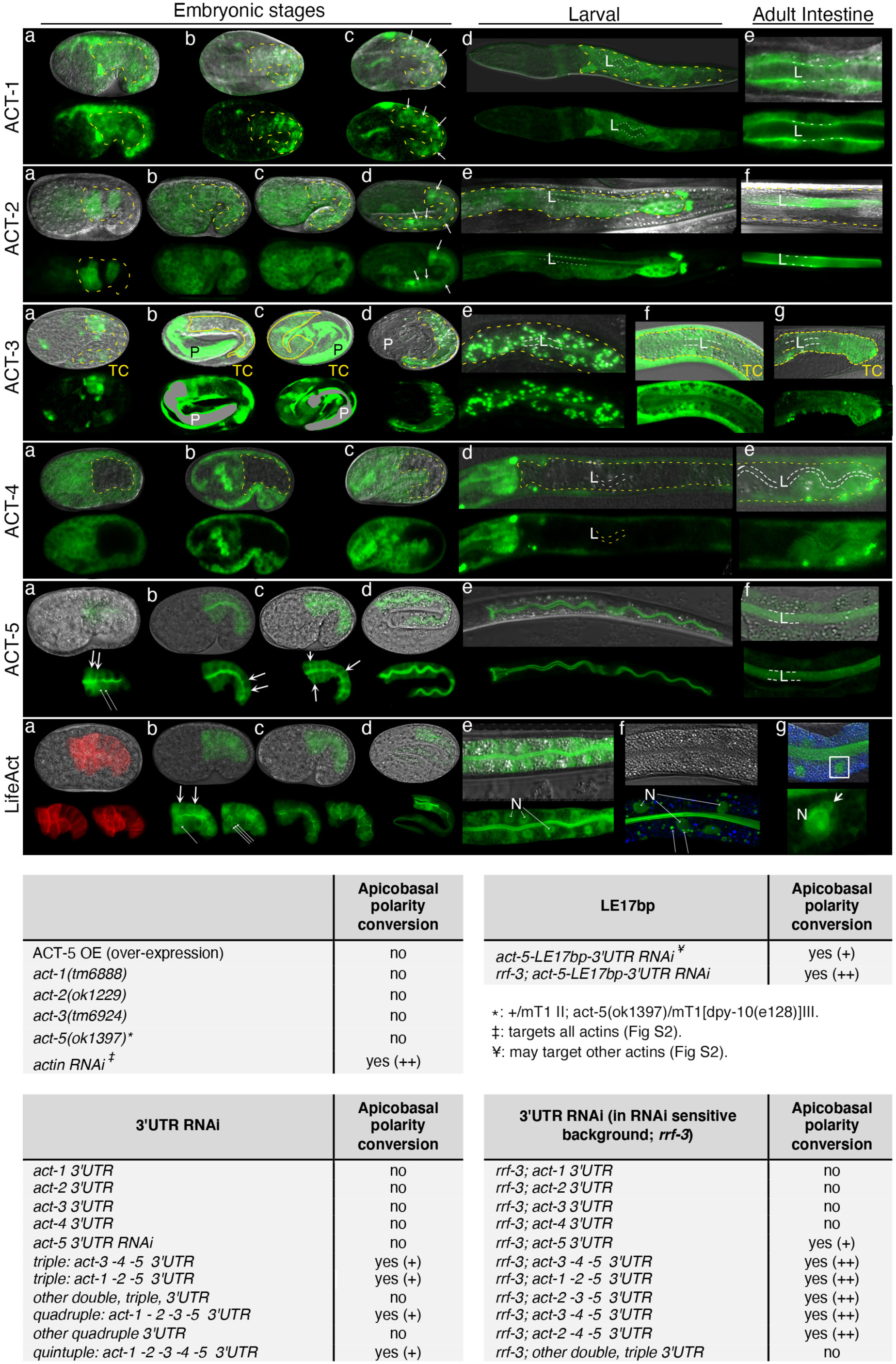
Multiple actin isoforms are expressed in the *C. elegans* intestine and function with ACT-5 in polarity. Note that ACT-1-4 expression is predominantly extra-intestinal (Fig.S5) and its intestinal expression is weak compared to ACT-5. **ACT-1::GFP:** (a-e) Comma, 2-fold, 3-fold, L1 and adult intestines with faint cytoplasmic (a-e), pan-membranous (apico-basolateral; b,c) and increasingly apical (d,e) ACT-1. Here and below, intestines are outlined by yellow dashes; arrows indicate basolateral membranes; L=lumen. **ACT-2::GFP:** (a-d) Embryonic stages with faint cytoplasmic and apico-basolateral intestinal ACT-2 (d), shifting to the apical membrane (larva in (e); adult in (f)). **ACT-3::GFP:** (a-c,f,g) A transcriptional *pact-3-gfp* plasmid (**TC**) is expressed in embryonic (a-c), larval (f) and adult (g) intestines. (d, e) A translational *pact-3-act-3-gfp* plasmid is expressed in vesicles and at the apical membrane in 3fold embryonic (d) and L1-larval (e) intestines (animals fail to survive beyond this stage*). For clarity, intestine is fully outlined by yellow line and pharynx (P) greyed out in b and c (intestine partially obscured by strong extra-intestinal expression). **ACT-4::GFP:** (a-d) No ACT-4 was detected in pre- and post-intercalation embryonic and early larval intestines. Faint cytoplasmic ACT-4 expression in the posterior intestine of older larvae (e). Location of lumen (L) indicated by dashed white line. **ACT-5::GFP:** (a-f) Shift from cytoplasmic/apico-basolateral (arrows; a-c) to definitive apical (d-f) ACT-5 location through intestinal intercalation and membrane polarization (a-d). Note occasional patches (long thin arrows) throughout embryonic development (a-d). **LifeACT::mCherry and LifeACT::GFP:** (a-g) ACT-1-, 2-, 3- and 5-similar LifeAct shift from cytoplasmic/apico-basolateral membrane (a-c) to apical membrane through intestinal intercalation and polarization (a-d). Note cytoplasmic mesh (a) and patches (b: two sections; long thin arrows) in embryos, and persistent cytoplasmic, peri-vesicular (long thin arrows in f) and peri-nuclear localization (N: nucleus, and emanating arrows) in larval (e), and adult intestine (f, g). Blue distinguishes autofluorescent lysosome related organelles (LROs). Confocal and confocal/Nomarski overlay images of full embryonic and partial larval and adult intestines are shown. See Methods for generation of fluorophore fusions, Fig.3S for schematics of intestinal development and polarized membrane expansion. **Table.** Summary of effects of single and combined mild perturbations of actin isoforms on intestinal polarity (related to STab.1; see Fig.S6 for comparative structural analysis of actin isoforms and Methods for rationale of 3’UTR approach). *The *pact-3-act-3-gfp* plasmid robustly and apparently specifically localizes to vesicular structures and to apical membranes of the developing intestine, a localization not compatible with survival (no transgenic lines could be generated). It can therefore not be excluded that ACT-3 transiently localizes to vesicles in the intestine (compare the occasional vesicular location of LifeAct). The ACT-3 protein is 100% identical to ACT-1 (Fig.S7). It is therefore also formally possible that the *act-1* promotor is weaker than the *act-3* promotor such that ACT-1 (below detection level), like ACT-3 (toxic as GFP fusion), also transiently localizes to vesicles.

**Figure 8:**
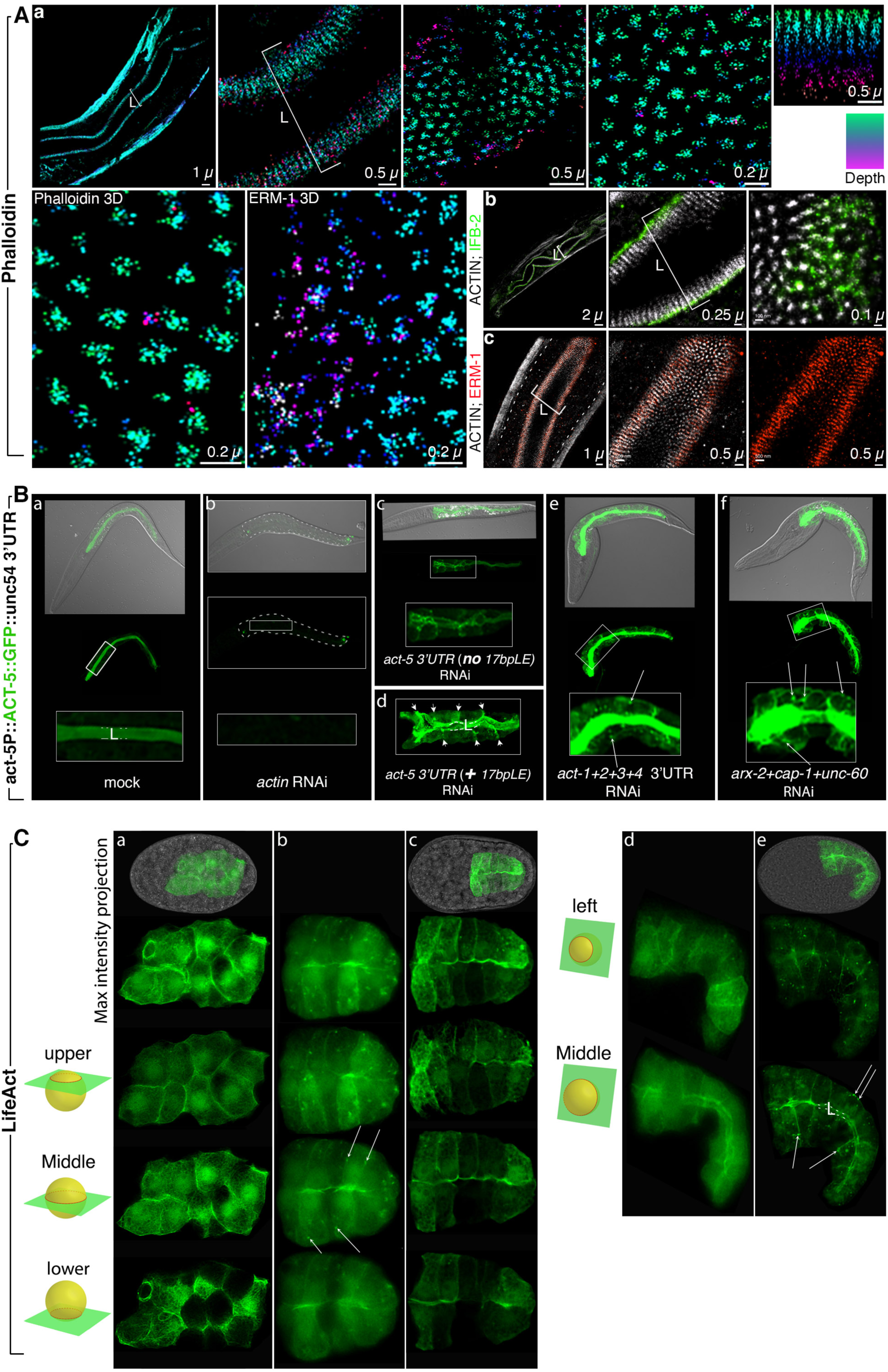
Actin determines its own polarity. **A: Stochastic optical reconstruction microscopy (STORM) imaging of the submembranous apical (lumenal) cytoskeleton of the L1-larval *C. elegans* intestine.** (a) 3D STORM (surface/lumen close = green; depth/cytoplasm close = purple) with Alexa Flour 647 phalloidin. Phalloidin staining of fixed worms was used to detect all actins and for best resolution (superior incorporation of phalloidin into actin filaments as compared to actin fluorophore fusions ^54^. Increasing magnification shown from left to right, sagittal sections through intestine and apical lining to the left, followed by 2 transverse and one sagittal section through microvilli; highest magnification in lower row resolves single actin filaments in microvilli. Note absence of circumferential actin belt in all images. (b-c) Double color STORM with Alexa Flour 647 phalloidin and Dy750- and Cy3-labeled nanobodies to IFB-2::GFP (b) and ERM-1::GFP (c), respectively. Note rooting of actin filaments in intermediate filament lattice (IFB-2) and full overlay of actin filaments and ERM-1. L=lumen. **B: Actin determines its own polarity (*act-5p*-ACT-5::GFP-*unc-54*-3’UTR).** (a) *act-5p*-ACT-5::GFP-*unc-54*-3’UTR is located at the apical domain (lumen) in the wild-type intestine and is efficiently removed by *actin* RNAi (b) that targets exonic sequence of *act-5* as well as other actin isoforms (Fig.S2b). (c-d) In contrast, *act-5-3’UTR* RNAi and *act-5LE17bp-3’UTR* RNAi, the former exclusively, the latter chiefly, targeting the *act-5* 3’UTR (Figs.S2b, S6a; missing in *act-5p-*ACT-5::GFP-*unc-54*-3’UTR), reveal that ACT-5/actin is required for its own polarized (apical) positioning (basolateral ACT-5 displacement indicated by arrows in d). (e) Quadruple *act-1-4* 3’UTR also displaces ACT-5 to the basolateral membrane and the cytoplasm (long arrows point to patches). (f) RNAi with the 3 bcAMs displaces ACT-5 to basolateral membranes, a filamentous cytoplasmic and cortex-associated structure and patches (long arrows). Note body morphology defects in b, e, f (elongation defects in all, ‘big head’ in e and f) but not in c, d (*act-5* specific depletion), indicating extra-intestinal RNAi effects of other isoforms and bcAMs and documenting efficiency of mild RNAi approach. Confocal and Nomarski/confocal overlay images of L1 larvae shown throughout, magnified areas indicated, intestine outlined by dashes in b, L=lumen. **C: A cortex-enriched cytoplasmic F-actin mesh recedes towards the apical domain during polarity establishment (LifeAct::GFP).** Top panels show confocal/Nomarski overlay images with corresponding confocal projections beneath (a, c, e). The 3D schematic spheres indicate focal planes of the corresponding confocal sections. (a) Pre-bean stage (pre-intercalation) embryonic intestine with dense cytoplasmic actin mesh, perinuclear actin, and actin enrichment along all membranes. Upper (dorsal) section shows that nuclei have not yet migrated towards the midline (future apical domain); middle section reveals that the actin mesh traverses the cytoplasm; lower (ventral) section shows effect of ventrally encroaching germ cells (black). Images are 1x deconvolved (2D blind; numerical aperture 1.4; refraction index 1.51; iteration 1; specimen thickness: thin; noise level: clear). (b, c) Late pre-bean stage (image in c 1x deconvolved). Actin mesh receding (compare a and c), less actin at basolateral membranes, more actin at midline (future apical domain), new actin patches (middle plane reveals that these are mostly located at or close to the cortex [long arrows]). Note apical translocation of nuclei at this stage of polarization ^23^. (d-e) 1.5-fold embryos (image in e 1x deconvolved). Note: further receding of cytoplasmic actin mesh towards the apical domain; pre-dominantly cortex-associated actin patches (long arrows in middle section) and lumen formation (L). F-actin continues to be enriched at the apical domain (lumen).

bcAM modeled branched actin networks can produce force without myosins and actin filament turnover (treadmilling) can generate vectorial dynamics that propels membranes forward, also documented in *C. elegans* ^31^. We next explored whether the cytoplasmic and/or cortex-based filamentous actin assembly itself, powered by bcAMs, might confer directionality to vesicles. An actin structure that moves towards the apical domain might also direct vesicles that can attach to this structure (e.g., via their previously identified apical polarity cues). Such a self-organizing vectorial actin structure should effect its own polarization. We used the ACT-5::GFP fusion missing the *act-5* 3’UTR to test this prediction (Fig.8B). Indeed, RNAi directed to the *act-5* 3’UTR (*act-5-3’UTR-* and *act-5-LE17bp-3’UTR* RNAi; Fig.2S) mislocalized apical ACT-5::GFP (no *act-5* 3’UTR; Fig.8Ba), effectively removed by *actin* RNAi (Fig.8Bb), to basolateral domains and the cytoplasm in larval intestines and increased its basolateral and cytoplasmic location in the intercalating embryonic intestine (Fig.8Bc-d; not shown; Fig.2S, 8 legends for details). Moreover, single, double, triple and quadruple 3’UTR RNAi with *act-1-4*, none able to induce polarity conversion on its own, mislocalized ACT-5::GFP to the basolateral domain and to cytoplasmic or cortex-associated patches (Fig.8Be; see Supplementary Results for: (1) the identification of ACT-1, -2, -3 and -4::GFP in the *C. elegans* intestine; (2) an ACT-5 similar developmental basolateral-to-apical polarity switch of ACT-1-3; (3) the non-redundant contribution of these actin isoforms to actin’s polarity function; Figs.7, S5). Finally, the combined depletion of UNC-60, ARX-2, and CAP-1 strongly displaced ACT-5::GFP to basolateral domains and to a filamentous structure in the cytoplasm (Fig.8Bf). We conclude that ACT-5, ACT-1-4, and the 3 bcAMs are non-redundantly required to promote actin’s own polarization to the apical domain during *de novo* polarized membrane biogenesis in the embryonic and larval epithelium. These results also reveal that ACT-1-4 are required to polarize ACT-5, consistent with these isoforms’ contribution to an ACT-5 containing dynamic actin structure that determines its own polarity.

To examine *in vivo* F-actin dynamics during membrane polarity establishment in the intercalating early-embryonic intestine, we used LifeAct as the most sensitive and actin isoform unbiased tracer of *in vivo* filament dynamics (see above, Figs.4, 7, 8C). LifeAct-GFP revealed a pan cortex-enriched cytoplasmic F-actin mesh in still dividing pre-intercalation intestinal cells that receded during intercalation, leaving transient patches along the cortex while becoming asymmetrically enriched at the nascent apical domain, before and during apical domain (lumen) biogenesis (Fig.8Ca-c). In postmitotic but still expanding late-embryonic and larval cells (E20 stage and beyond), this mesh continued to recede while F-actin increased at its destination, the apical domain (see Figs.7, 8Cd-e and legend for details). Together, these findings were consistent with a self-organizing polymeric F-actin structure that asymmetrically shifts across the cytoplasm and/or along the cortex towards the apical membrane domain during the concomitant proce ss of this domain’s biogenesis and positioning.

A positional F-actin shift from the basolateral to the apical domain, rather than reflecting active movement, could be the consequence of actin’s successive exclusion from the basolateral domain during membrane polarity establishment (with or without new F-actin assembly at the apical domain). Domain exclusion of polarity cues, e.g., the PARs, is considered a principle mechanism of specifying membrane polarity in the epithelial tissue context (see Introduction), and actin’s polarity shift, rather than initiating the membrane polarity change, might be a consequence of it. Such a scenario would be impossible to exclude even by time lapse imaging. To directly test the hypothesis of a dynamic F-actin structure that assumes apical directionality, we generated photo-activatable LifeAct::PA-GFP and photo-convertable LifeAct::Dendra2 fusions, directed them to the intestine and examined the translocation of activated GFP and converted Dendra2 in single intestinal cells and within the whole intestine during *in vivo* polarity establishment in real time (Figs.9, S9; legends and Methods for details). Activated LifeAct::PA-GFP and converted Life-ACT::Dendra2, both detecting only the initially activated/converted, but no newly generated, F-actin, was indeed found to translocate from its basolateral membrane location through the cytoplasm and/or along the cortex to the nascent apical domain of intestinal cells, at the time of polarity establishment and during the ongoing process of *de novo* polarized membrane biogenesis in the developing embryonic epithelium (Figs. 9, S9; movie 1&2).

**Figure 9.**
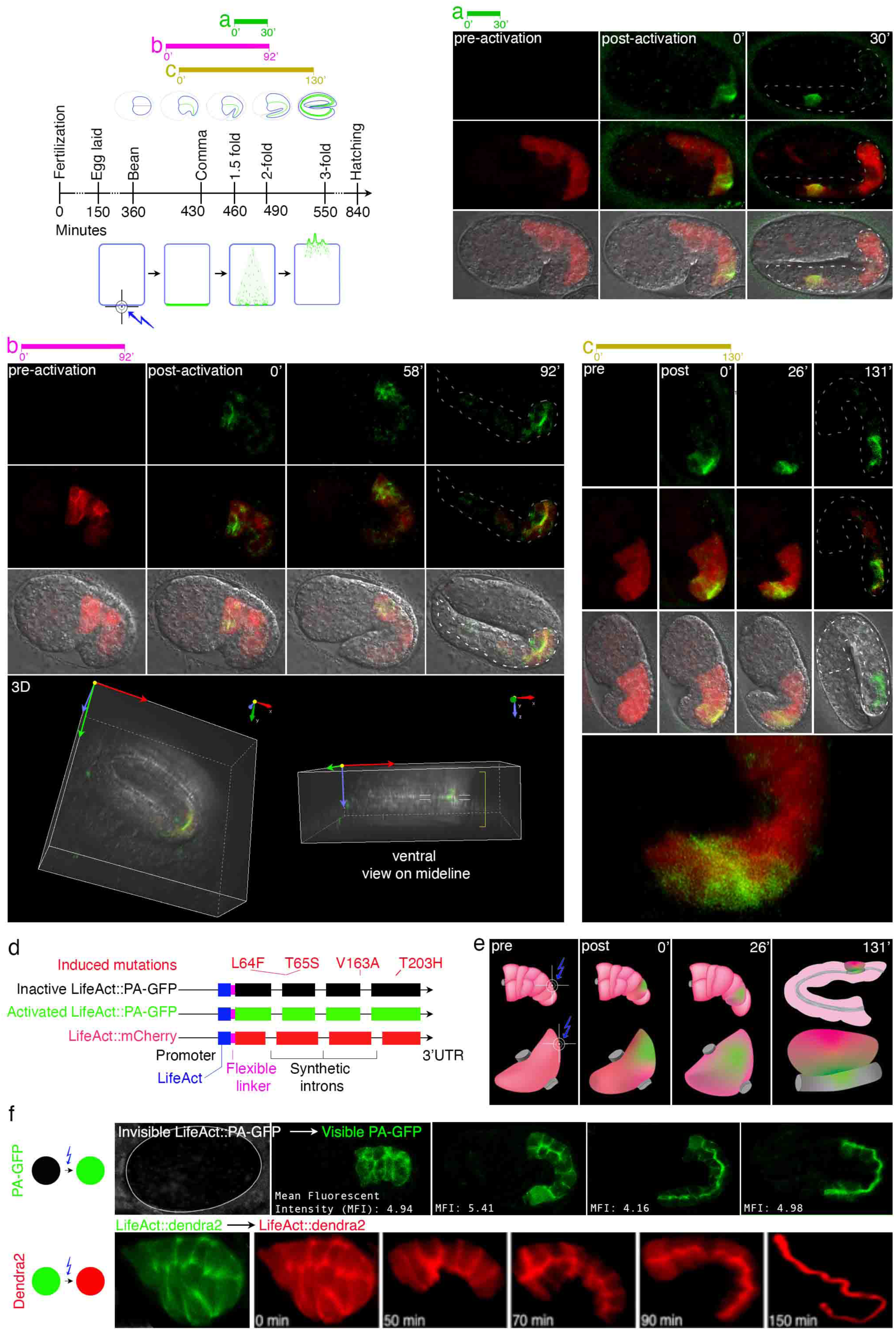
Tracking the transcytotic F-actin shift from the basolateral to the apical domain with time lapse imaging of photoactivated and photoconverted LifeAct. Upper left panel shows timing of image recording relative to timing of embryonic development. a, b, and c horizontal timelines correlate to a, b, c of the 3 following sets of images (see Fig.S8 for additional timelines using photoconversion with LifeAct::Dendra2). Photo-activation schematic: a basal (or lateral) membrane of one single cell is targeted (blue lightening sign) and LifeAct::PA-GFP tracked. Direction of GFP shift is presented as arrow, GFP quantity remains constant (newly generated LifeAct is invisible). (a-c) To allow for precise photoactivation of LifeAct::PA-GFP (not visible before activation) at the basal or lateral membrane domain of one single cell, animals were also supplied with LifeAct::mCherry that outlines intestinal cells including basolateral membranes. (a) Photoactivation in a 1.5-fold embryo (post intercalation intestine). Basolateral membrane angle of one posterior intestinal cell is targeted using LifeAct::mCherry (red) as guide (green emission of PA-GFP at 0 minute). 30 minutes later the PA-GFP is retrieved at the apical domain, demonstrating translocation. (b) Photoactivation at late bean stage (during intestinal intercalation). Lateral membranes of two anterior cells (E3L and R) are photoactivated (Fig.S3 for E cell development and intercalation). PA-GFP is visible in cytoplasmic or possibly cortex associated patches after 58 minutes and arrives at the apical membrane at 92 minutes. Rotated 3D images below show GFP location at the midline (future lumen). (c) Photoactivation at comma stage. Basal portion of basolateral membrane of one posterior intestinal cell is targeted. Apical directionality of PA-GFP translocation visible after 26 minutes (enlarged beneath) and traced to the apical membrane in the fully formed intestine of the 3-fold embryo at 131 minutes. (d) Map of LifeAct constructs. Four mutations were introduced into the GFP to make it photo-activatable (Methods). LifeAct (shown in blue; only 17 amino acids). A flexible linker (7 amino acids) keeps fluorophore in distance from protein. Synthetic introns are optimized for brighter expression in *C. elegans*. (e) Schematic tracking events shown in c to clarify LifeAct translocation in single intestinal cell and within the organ (embryo shown in c rotated 90° counterclockwise). Targeted area shown by blue lightening sign. LifeAct::PA-GFP dynamics (green) are followed from basolateral to apical membrane (lumen) in one cell (lower panel) and within the developing intestine (upper panel; lumen = green). (f) LifeAct photoactivation and photoconversion of whole intestines and tracking of fluorophores throughout intestinal development, from pre-intercalation (left) to full organ development in 3- fold embryo (right). Note that mean fluorescence intensity remains constant throughout development after PA-GFP activation. See Fig.S9 for LifeAct::Dendra2 conversion and single-cell tracking. Confocal and confocal/Nomarski overlay images are shown throughout. Intestines are outlined by dashed lines in a - c.

## DISCUSSION

We here identify branched-chain actin dynamics as an epithelial polarity cue in the *C. elegans* intestine. Neither actin itself, a key polarity cue in budding yeast, nor the components of its branched chain assembly are currently considered *bona fide* polarity cues in epithelia, despite their functions in multiple aspects of cell asymmetries^1^. In the *C. elegans* intestine, actin has been considered altogether dispensable for polarity^15^. The independent identification of the 3 principle components of the branched-chain actin machinery (UNC-60/cofilin, ARX-2/Arp2/3 component, CAP-1 capping – designated branched chain actin modulators/bcAMs) among a limited number of molecules identified in genome-scale screens by the same unique loss-of-function polarity phenotype, strongly supports the physiological relevance of branched-chain actin dynamics for this function.

Several lines of evidence suggest that the here identified regulatory function of bcAMs/actin in membrane polarity is mediated by their effect on the apical directionality of anterograde vesicle trajectories at the time of membrane polarity establishment, rather than on their canonical functions, some with effects on polarity (e.g., apical cortex and junction modeling, lumenogenesis, polarized trafficking and directional cell migration ^18, 19, 32^: (1) the bcAM/actin-dependent polarity phenotype copies the polarity phenotype induced by interference with vesicle-based polarity cues previously shown to direct biosynthetic trafficking to the apical domain^7^; (2) bcAMs/actin functionally interact with these vesicle-based polarity cues in polarity; (3) all components of branched chain actin dynamics are required to direct pre- and post-Golgi vesicles to the apical domain during *de novo* polarized membrane biogenesis; (4) both systems (actin dynamics and trafficking) operate in polarity upstream of apical PARs; (5) all components of the system (bcAMs; several actin isoforms; actin itself; vesicles) shift location through the cytoplasm/along the cortex from the basolateral to the apical domain at the time of polarity establishment; (6) a polymeric, bcAM-driven F-actin structure physically translocates towards the future apical domain during polarity establishment. We therefore propose that bcAMs and actin function in epithelial polarity by constraining the essential biosynthetic-secretory trafficking pathway, required to supply all sides of the membrane, from the ER towards the apical domain, a process we suggest establishes membrane polarity and maintains it as long as new polarized membrane is synthesized. Branched chain actin dynamics could provide the underlying cytoskeletal structure and thus the proposed mechanism for a regulated directional change of vesicular trafficking that is independent of polarized cargo sorting to cognate recognition sites at already polarized target membrane domains^7^.

Our findings thus support a polarity model where the insertion of the apical domain into the growing membrane partitions its apicobasal membrane domains and recruits the membrane-based polarity cues (proposed model^7^, rather than a model where membrane-based apicobasal polarity cues recruit polarized membrane components by directing intracellular processes such as vesicular trafficking (current model^1^. Although the proposed model suggests a different ordering of events, it does not challenge any components of the current model. Beyond their function in providing platforms for the recruitment of polarized membrane components, the membrane-based core polarity cues (e.g., apical PARs and junctions), are well placed to subsequently or concomitantly (with the directional delivery of apical membrane) secure: (1) the growing apical domain (e.g., via the apicolateral assembly of junctions); (2) the vesicle trajectories that expand the apical domain (e.g., by tethering them to the apical membrane); and (3) the established membrane domains - via their intricate signaling network of mutual apical versus basolateral inhibition: ^33–38 39–43^. We suggest that the proposed polarity model, like all its identified, highly conserved molecular components, is conserved between species, epithelia and other tissues, as is the here characterized epithelial polarity phenotype (e.g., in the MDCK cyst model^9^) and the increasingly appreciated role of intracellular vesicular trafficking in apical/anterior domain positioning and maintenance in diverse tissues and species^8, 9, 12, 13, 44, 45^. The model has precedence in budding yeast, where all its critical components (actin, secretion) were the first to be identified in screens for polarized growth ^46^, and where actin’s function as polarity cue is chiefly mediated by directing the secretory pathway to the bud site ^47^. All 3 components of yeast bcAMs function in the regulation of cortical actin patches, critically involved in polarized cell division ^48^. Moreover, the asymmetric insertion of the apical domain also initiates membrane polarization in the mouse embryo, an event similarly characterized by the apical positioning of PARs and ERM/ezrin ^49^. Polarization processes from yeast to humans provide membrane-based signals other than already polarized membrane domains that could reorient vesicle trajectories in epithelia (e.g., the midbody that positions the apical domain in MDCK cysts and moves to the future apical domain in the polarizing *C. elegans* intestine ^14, 50–52^. A self-organizing cytoskeletal system is highly flexible and amenable to be directionally regulated in various ways in the epithelial tissue context, cell-autonomously or non-autonomously, transcriptionally or post-transcriptionally, developmentally or dependent on the regulation of polarized cell division or migration (see discussion in ^7^; see Supplementary Discussion for review of distinct types of dynamic actin assemblies that could serve as candidates for the here described transcytotic polymeric F-actin shift that occurs during the process of membrane polarity establishment).

## Acknowledgements

Strains and reagents were kindly provided by:

Barth D. Grant (Rutgers University, Piscataway, New Jersey, USA),

Kenneth Kemphues (Cornell University, Ithaca, New York, USA),

Keith Nehrke (University of Rochester Medical Center, Rochester, New York, USA),

Shoichiro Ono (Emory University School of Medicine, USA),

Erwin J Peterman (Vrije Universiteit, Amsterdam, Netherlands)

Jonathan Pettitt (Institute of Medical Sciences, Foresterhill, Aberdeen, UK),

Scott A Rifkin ^55^ (University of California San Diego),

Christian E. Rocheleau (McGill University, Montreal, Quebec, Canada),

David Sherwood (Duke University, USA),

TEM work was performed at the Center for Systems Biology/Program in Membrane Biology (Massachusetts General Hospital/MGH, Boston, USA).

The *C. elegans* Knockout Consortium^56^

The *C. elegans* National Bioresource Project of Japan (NBRP) The Caenorhabditis Genetics Center.

Y. Kohara (National Institute of Genetics, Mishima, Japan)

We thank David Hall (Albert-Einstein College of Medicine, New York, USA) for discussion of ultrastructural images, John Fleming and Frank Solomon for critical reading of the manuscript, and Howard Weinstein and Ronald Kleinman (MGFfC, Harvard Medical School, Boston, USA) for ongoing support.

## Supplementary Materials

Materials and Methods Figs. S1 to S8

Caption for Movie S1 and S2

Tables S1 and S2 (please see Excel files) Supplementary Results

Supplementary Discussion References

Movie S1 Movie S2

