## Supplementary Materials for "A transcytotic actin shift polarizes vesicle trajectories and partitions apicobasal epithelial membrane domains"

**This PDF file includes:**

Materials and Methods

Figs. S1 to S8

Caption for Movie S1 and S2

Tables S1 and S2 (See Excel file).

Supplementary Results

Supplementary Discussion

References

**Other Supplementary Materials for this manuscript include the following:**

Movie S1

Movie S2

#### Materials and Methods

##### Experimental model
An introduction to the *in vivo* analysis of polarized membrane biogenesis in the *C. elegans* intestine, with an extended Methods section and a visual demonstration of the techniques used in this study is provided in[1, 2].

##### *C. elegans* Strains, Culture Conditions, Genetics
Wild type (N2 Bristol) and mutant *C. elegans* strains were cultured and genetic crosses performed using standard methods[3]. Worms were generally maintained at 20-22°C (unless otherwise noted) on Nematode Growth Medium (NGM) plates seeded with *E. Coli* OP50[4]. The list of strains used in this study is provided in Table S2.

##### RNA interference
###### General
Methods have been previously described[5]. Briefly, RNAi was carried out by feeding worms *E. coli* HT115 (DE3), producing double stranded/ds RNA of the gene of interest, as previously described.[6, 7]To avoid OP50 contamination, prior to seeding worms onto RNAi plates, animals were washed three times with M9[3] and/or swirled on the plate in drops of carbenicillin solution (500mg/ml). For standard RNAi, bacterial feeding clones were inoculated from LB plates into 1ml LB liquid medium containing 50μg/ml ampicillin and incubated for 8-18 hours at 37℃. 200μl cultured RNAi bacteria were seeded onto agar plates supplemented with 2mM IPTG and 25μg/ml carbenicillin. dsRNA was induced at room temperature for at least 6 hours before picking 4 - 6 L4 larvae onto each RNAi plate. Most bacterial clones were derived from the Ahringer genome-wide RNAi feeding library (J. Ahringer, Welcome Trust/Cancer-Research-UK-Gurdon-Institute, Cambridge, UK). The integrity of all RNAi clones was verified by sequencing.

###### Scaled intensity RNAi approach
**Rationale.** The purpose of this study was to generate informative loss-of-function phenotypes for highly pleiotropic genes (actin and actin modulators), in order to distinguish their specific function in polarized membrane biogenesis from various other functions. All genes are sterile, maternal-effect and embryonic/early-larval lethal. RNAi, rather than tissue-specific protein degradation strategies, was therefore determined as the method of choice. It is easily titrated to produce a range of mild-to-moderately severe phenotypes and it effectively targets maternal RNA (limiting the use of germline mutants due to the requirement of balancers that provide maternal product). Severe reduction- or full loss-of-function phenotypes, even if tissue-specific (e.g., directed to the intestine), were found to be less informative due to the disruption of basic cellular processes, such as cellular morphogenesis and vesicular trafficking (see text).
**Procedures.** Mild-to-moderate RNAi conditions were empirically determined for any given gene by: modulating IPTG concentrations in RNAi plates; diluting the RNAi clone of interest with different amounts of mock bacteria (no vector or vector with irrelevant gene); varying the developmental stage (L2 to adult) of the parental strain in which RNAi was induced; using the RNAi-sensitive strain *rrf-3(pk1426)*.[2] Conditions were furthermore varied to achieve interference at different temperatures (15°C, 22.5°C, and 25°C) and different time points during embryonic

and/or later stages of development, e.g., RNAi was induced: in parents (evaluating the F1 progeny; standard parental RNAi); in larvae (evaluating the same generation; conditional larval RNAi); or in adults. Conditional larval RNAi was generally carried out by bleaching 30-50 gravid adults in one drop bleaching solution (a 1:4 mix of 10M NaOH and household sodium hypochlorite) on the edge of an RNAi plate and allowing hatched larvae to crawl to the bacterial lawn (see[2] for details). Conditional RNAi was also induced at different stages of development by transferring eggs, L1, L2, L3, or L4 larvae (larval RNAi), or adult animals to RNAi plates and scoring the same generation (not their progeny, as in standard, parentally induced RNAi) over time. Appropriate controls were added to ensure that RNAi was effective when induced at later time points during development or in adults (e.g., by using *gfp* RNAi on a GFP expressing strain). All experiments were repeated three or more times for each dataset.

**3'UTR RNAi for actin isoforms**

**Rationale.** The genetic analysis of actins has been impeded by the large families of multi-functionally essential, often maternally required, almost identical and redundant actin isoforms, whose stoichiometries are furthermore critical for filament assembly. Loss-of-function analyses typically fail to produce effects if targeting single isoforms. If effective, they induce early lethality when used globally, or severe cellular morphogenesis defects if used in a tissue-specific manner (e.g., via protein degradation strategies). Mutational and gain-of-function approaches have historically proven more effective, often induce dominant negative, but may also induce neomorphic changes (e.g., a dominant *act-2* germline mutation, but not an *act-2* germline deletion, produces embryonic morphogenesis defects in *C. elegans*[8]). We trialed a 3'UTR directed RNAi approach expected to induce mild phenotypes for the reasons outlined above (scaled intensity RNAi approach: to avoid cellular morphogenesis and intracellular trafficking defects.) Moreover, it was effective as combinatorial approach, up to quintuple combinations, facilitating the analysis of redundancies between isoforms.

**Procedures.**

As shown in Figure S6, the 3'UTR of each gene were selected right after the stop codon and, using the Clone mapper tool,[9] the "Off-Target" regions (shared with other actins) were excluded. The desired (specific) 3'UTR region was generated by PCR and inserted (Gibson method) into the multiple cloning site of the L4440 backbone (a modified version of the pBlueScript plasmid) [10] and sequence validated. The plasmid then was transfected into the *Escherichia coli* strain HT115 (DE3)—a strain carrying a defective RNase III and an isopropylthiogalactoside (IPTG)-inducible T7 polymerase gene[11].

Combined 3'UTR or other RNAis were prepared by mixing the desired proportional RNAi liquid volumes of each desired combination (for example, 1-1 volumes, or 50% each) prior to seeding the RNAi clones onto agar plates supplemented with IPTG and carbenicillin (see above). Appropriate controls (corresponding dilutions with mock RNAi bacteria) were included for combinatorial RNAis in all experiments.

**Genetic Interactions**

**Rationale.** Most genes examined in this study are maternal-effect and early lethal genes, prohibiting the use of null mutants in genetic interaction experiments. Moreover: (1) full loss of gene function was not desired, since expected to mask specific effects on membrane polarity (see above, scaled intensity RNAi approach); (2) only RNAi will remove maternal product that is delivered by the obligate balancers for germline null mutants; (3) enhancement was expected for

all combinations tested in this study. We therefore generated very mild reduction-of-function conditions by using temperature sensitive mutants, heterozygous and very mild RNAi conditions (see above, RNAi).
**Procedures.** See above, Genetics and RNAi.

**Analysis of *de novo* polarized membrane biogenesis in single postmitotic cells of the expanding intestine**
See[5] for the postmitotic larval intestine as a model for the *in vivo* analysis of *de novo* polarized membrane biogenesis. This model allows the separation of *de novo* polarized membrane biogenesis from confounding effects of polarized cell division and migration, a difficulty for the analysis of polarity establishment (typically investigated in proliferating and moving cells). In addition, polarity conversion conditionally induced during larval membrane expansion in postmitotic cells identifies direct effects on polarized membrane biogenesis by excluding the possibility that the polarity defect is a consequence of anteceding polarized tissue morphogenesis defects. Note that these effects can be distinguished from membrane polarity maintenance in non-expanding mature epithelia[5]. See Fig.S3 for net membrane addition in the late embryonic and larval *C. elegans* intestine.

**Measurement of apical membrane length in embryos and larvae**
The length of the apical/lumenal membrane of the intestine at all stages of development from the embryo to the adult was measured from confocal images, using the "Annotation and Measurement tool: polyline length tool" provided in the NIS-Elements AR software. The results were computed by a simple statistics table and exported into an Excel sheet. Triplicates of 15 animals each were examined. Image pixels: 512 x 512. Objectives used: for embryo and L1: 60x. for L2-3 larvae: 20x; for L4-larvae and adults:10x.

**DsRed feeding**
Methods have been previously described.[5] DsRed HT115 RNAi bacteria were generated with a DsRed expressing plasmid to produce a faint red color. Animals were fed on plates containing RNAi bacteria targeting secretory pathway components and control RNAi bacteria (no vector) for 2 days. At least 70 animals were transferred to plates containing a 1:1 mixture of gene-specific- and of DsRed containing RNAi bacteria, at least 15 hours before evaluation.

**Fluorescent fusion proteins**
All strains carrying fluorescently labeled fusion proteins are described in Table S2. The subcellular localization of most membrane and junction markers used in this study was previously confirmed by us and others by different labeling procedures, including distinct transgenes, antibodies, germline knock-ins, chemical staining; and by their ability to rescue the corresponding mutant phenotype (see text and[1, 2]).

**Rationale for generating exogenously tagged proteins.** All fusion proteins expected to directly affect actin filament assembly were exogenously rather than endogenously tagged and directed to the intestine to: (1) avoid interference with the pleiotropic functions of these essential components of filament assembly by changes at their germline loci (i*n vivo* actin filament assembly and dynamics are sensitive to changes in stoichiometries between actin and actin-binding molecules; see genetic interactions, Figs.2, S2); (2) support a high-resolution subcellular analysis of these ubiquitously expressed molecules in the developing intestine; (3) achieve optimal expression levels; (4) the purpose of the analysis was not the new identification of the gene-specific expression pattern, but the analysis of subcellular changes in localization during intestinal polarity establishment. All transgenic lines were generated with low copy number insertions and other safeguards against overexpression-induced artifacts.[1,2] Figs.4 and S4 also document overlap of transgene-derived and endogenous ARX-2::GFP and RFP (note vastly superior resolution achieved by the former). All genes examined in this study have previously been shown to be expressed in the intestine (except for the actin isoforms).

**Procedures.** To examine their subcellular localization in the early embryonic intestine, genes of interests were expressed from the intestine-specific *elt-2* promoter (promoter lengths vary between 600 and 5000 bp) at their 5′ ends and cloned in frame with GFP or tagRFP either at their 5′ or 3′ ends, using standard cloning procedures and Gibson Assembly cloning techniques[12]. We generated all bcAMs *(unc-60, arx-2, cap-1)* as both N-terminal and C- terminal fluorescent fusion proteins. The promoters (either *elt-2p* or the endogenous promotor, see text) were amplified by PCR from wild-type genomic DNA. cDNAs were PCR amplified from their respective cDNA plasmid clones (gifts from Y. Kohara, National Institute of Genetics, Mishima, Japan). GFP and tagRFP coding DNA fragments were PCR amplified from the plasmids ppD95.75 and pPD284, respectively (Fire et al., 1990; Addgene). *unc-54* , *elt-2*, or the endogenous 3′ UTR were used as indicated (see text). Plasmids were either cloned (as above) and/or generated using the PCR stitching method[13]; e.g. ACT-2::GFP was generated in both ways). All recombinant plasmids and PCR-stitched generated chimeric DNAs were sequenced prior to microinjection. To generate transgenic lines with extrachromosomal arrays, DNAs were injected into wild-type, *unc-119*, or mutant *C. elegans* gonads for germline transformation, with or without the *rol-6* marker or *unc-119+* rescuing construct, using standard microinjection techniques.[14, 15]

**Specific fusion proteins.** Also see[5]. **ERM-1::GFP.** The low copy number *erm-1p::erm-1::gfp* transgene, previously extensively characterized and shown to be devoid of any phenotypic effects, is suited for its use as apical domain identity marker since it avoids any disturbance of the *erm-1* germline locus that might interfere with apical domain biogenesis [7, 16-18]. ERM-1's subcellular localization has been previously confirmed by various independent transgenic strains (high and low copy number, labeled with different fluorophores), and by an ERM-1::GFP CRISPR knock-in [16, 17, 19]. All **vesicle-based fluorescent fusion proteins** are translational fusions driven to the intestine by the *vha-6* promotor to allow for the subcellular positional analysis of these ubiquitously expressed molecules (most generously provided by Barth Grant). Two strains of **ACT-3::GFP** were used, one translational fluorophore fusion, and one transcriptional fusion (see Fig.7 legend, Table S2). **PAR-3::mCherry, PAR-6::GFP and GFP::PKC-3** are CRISPR knock-ins (generously provided by Kenneth Kemphues). **LifeAct::GFP**. We found that the small exogenous F-actin-binding LifeAct combines inertness and sensitivity superior to endogenous actin-binding sites, such as the *Drosophila* MOE or *C. elegans* ERM-1 C-termini, respectively). LifeAct strains were backcrossed 4x and then integrated using UV irradiation to establish nonmosaic transgenic lines.[20]

**Dendra2 and PA-GFP strains.** A pFG102 vector was constructed, which contains 566 bp of *elt-2* promoter followed by LifeAct and Dendra2 or PA-GFP, the *elt-2* 3'UTR and a kanamycin selection marker. **Photoconvertible Dendra2** was PCR extracted from pEG412 from Addgene, which was sequence optimized for worm expression (three synthetic introns were inserted [21]), gel extracted, DPNI treated and replaced for the GFP in pFG102, using Gibson cloning technology. **Photoactivatable GFP (PA-GFP):** Initially described by Lippincott-

Schwartz [22] PA-GFP was used by Mijalkovic et al. in *C. elegans* [23]. We engineered our own PA-GFP using Genewiz DNA block and Gibson Cloning and introduced all four known PA-GFP mutations (L64F, T65S, V163A and T203H) into the Fire vector pPD95.75 GFP. Constructs were microinjected and transgenic strains generated.

**Immunohistochemistry**

Methods have been previously described[5]. Briefly, L1 larvae were collected in M9 medium[3] onto slides coated with 0.1-0.2% poly-L-lysine (Sigma, P5899), covered with overhanging coverslips, and then permeabilized by flash freezing in liquid nitrogen and subsequent flicking off the coverslip[2]. Fixation was performed by sequential incubation in methanol and acetone at -20℃. Immunofluorescent staining was carried out as described (procedures are demonstrated in[2]). For MH33 staining, slides were exposed to the first antibody (1:10 dilution) overnight at 4°C, washed and then exposed to the secondary antibody for 1 hour at room temperature. Permount (Fisher, SP15-100) was used as cover-slide mounting medium.

**Confocal and dissecting microscopy**

Methods have been previously described[5]. Briefly, differential interference contrast (DIC, Nomarski) and confocal images were acquired using a Nikon Eclipse-Ti inverted microscope equipped with C2 confocal system. All confocal images were obtained with a 63× objective. Exposure to fluorescent light was minimized to avoid bleaching, and images were obtained within minutes of mounting. Images were captured as single sections or a series of sections along the z axis with differing thicknesses (generally 0.1 to 1.0 μM). For multichannel images, individual channel intensity was adjusted, and the samples were scanned sequentially to exclude the possibility of bleed-through between channels. Confocal imaging parameters, such as pinhole size and LASER intensity, were empirically determined based on fluorophore intensity and experimental setting (avoiding phototoxicity, photobleaching, bleed-through). Deconvolution software was only used in Fig.8, as indicated, and images were not further edited except for adjustment of brightness and contrast (Adobe Photoshop).

**Fluorescence intensity measurement**

Using ImageJ[24] software, the entire image intensity was measured. Since all the PA-GFP and Dendra2 images were taken with the exact same criteria and settings on the same day (e.g.,, laser intensity, gain, pixels, etc.), background noise subtractions were not performed. Results were reported as Integrated Density. Triplicates of 15 animals each were analyzed.

**Transmission Electron Microscopy (TEM)**

Methods have been previously described[5]. Briefly, larvae were washed off in standard M9 medium[3] and collected into 1.5ml Eppendorf tubes. They were then fixed in 2.5% glutaraldehyde, 1.0% paraformaldehyde in 0.05M sodium cacodylate buffer (pH 7.4) plus 3.0% sucrose. Prior to fixation, the cuticles were 'nicked' with a razor blade in a drop of fixative under a dissecting microscope to allow the fixative to penetrate. After an initial 2-hour fixation at room temperature, the specimens were transferred into fresh fixative and stored overnight at 4°C. Specimens were rinsed several times in 0.1M cacodylate buffer, then post-fixed in 1.0% osmium tetroxide in 0.1M cacodylate buffer for 2 hours on ice. After post fixation, specimens were rinsed several times in 0.1M cacodylate buffer, then embedded in 2.0% agarose in PBS for ease of handling. The agarose blocks were dehydrated through a graded series of ethanol to 100%, dehydrated briefly in 100% propylene oxide and pre-infiltrated overnight on a rocker in a 1:1 mixture of propylene oxide:Eponate resin (Ted Pella, Redding, CA). The following day, the agarose blocks were infiltrated in 100% Eponate resin for several hours, then embedded in flat molds in fresh Eponate resin and allowed to polymerize a minimum of 24 hours at 60°C. Thin sections were cut on a Leica UC7 ultramicrotome and collected on formvar-coated grids, post-stained with uranyl acetate and Reynold's lead citrate, and viewed in a JEOL 1011 TEM at 80 kV equipped with an AMT digital imaging system (Advanced Microscopy Techniques, Danvers, MA).

**Stochastic Optical Reconstruction Microscopy (STORM)**
Coverslips rather than glass slides (length 50 mm, thickness 0.16; Fisherbrand, 12-544-E) were used to mount the worms. 30 µl of 0.01% Poly-L-lysine (Sigma, p5899) was added per two coverslips, sandwiched, separated, and air dried for 30 minutes. 5-10ul of 1xPBS was added and 50-100 animals mounted onto the slide (in 1xPBS). The second coverslip was added on top and with gentle pressure all animals were flattened. The sandwich was put on a -80°C metal block (using liquid nitrogen) and frozen for 5 minutes. Once frozen, one coverslip was flicked off. **Fixation and permeabilization**: All steps on ice. 5 µl 3% glutaraldehyde (Ricca, R3281800, 1.75 µl 10% triton, 43.75 µl CS buffer (10mM MES pH 6.1, 150mM NaCl, 5mM EGTA, 5mM glucose, 5mM MgCls) was applied for 2 minutes, followed by 30 µl 3% glutaraldehyde, 15 µl CS buffer applied for 10 minutes, followed by freshly made 0.1% NaBH4 (Sigma, 213462) in 1xPBS for 7 minutes. Sample(s) were washed three times using 1xPBS. The desired antibody was applied overnight at 4°C as following: Antibody labeling for actin: Alexa Flour 647 Phalloidin (Fisherbrand, A22287) conjugated 9:100 in 1xPBS; other antibodies used: anti-GFP nanobody AL647 (Alexa Flour 647) conjugated 1:100 in blocking solution, IFB-2 primary antibody MH33 1:10 and secondary donkey anti-mouse AL647. Animals were imaged in PBS buffer containing 100 mM cysteamine (Sigma), 5% glucose (Sigma), 0.8 mg/mL glucose oxidase (Sigma), and 40 µg/mL catalase (Roche Applied Science).[25]

**Photoconversion (of Dendra2) Procedure:** Mounting: early embryonic, bean, comma or 1.5-fold stage animals were selected and transferred onto agarose pads. To avoid hypoxia, we created a square-shaped 3mm x 3mm 5% noble agar pad that has enough O2 ventilation and mounted 4-5 transgenic embryos onto a 1 µl drop of M9 on the pad, covered with standard glass coverslip. **Photoconversion:** Using a laser scanning confocal microscope equipped with a 63X objective, prior to photoconversion, a z stack image was taken using FITC and TRITC channels and customizing parameters to avoid bleed-through between the channels (TRITC {Laser Wavelength}: 488.0, [26]: 3.2, {PMT HV}: 70, {PMT Offset}, and FITC {Laser Wavelength}: 561.0, [26]: 6.0, {PMT HV}: 78, {PMT Offset}: -15, and TD {PMT HV}: 89, {PMT Offset}: 0). Using ROI tab, the region of interest was selected and stimulation (UV 405nm, C2plus stimulation as 0.36. ND stimulation: 1 sec, 3 loop) was applied. Another post conversion z-stack image was taken at minute 0, followed by images with desired minute intervals as the embryos develop.

**Photoactivatable GFP (PA-GFP) Procedure**: Mounting as above (photoconversion). **Photoactivation**: Using a laser scanning confocal microscope equipped with a 63X objective, an image of the worm was acquired (as described above for Dendra2 photoconversion). Next, using 405nm Ultraviolet (UV) Laser, a region of interest in the embryonic intestine was photoactivated using a customizable region of interest (ROI) bleaching tool within the laser scanning confocal acquisition software. For our studies, optimal photoactivation was achieved using the 405nm laser at 10% power. Worms were imaged immediately following photoactivation. It is important to ensure that the image acquisition setting (e.g., exposure, gain, laser power, binning, etc.) are maintained during intervals of imaging sessions or experiments.

**Statistics**

Statistical analyses were performed by GraphPad Prism 9.3.0 (Mac) software. All values are mean $\pm$ SEM of three or more independent experimental data sets. P values were calculated by student's two-tailed *t-test*. n (sample size) is indicated in the text and in the figure legends where necessary. $^{*}p < 0.05$, $^{**}p < 0.01$, $^{***}p < 0.001$.

**Software used:** Clone Mapper[9], Transfac T-coffee[27], ImageJ[24], Clustal Omega[25].

Fig 1S

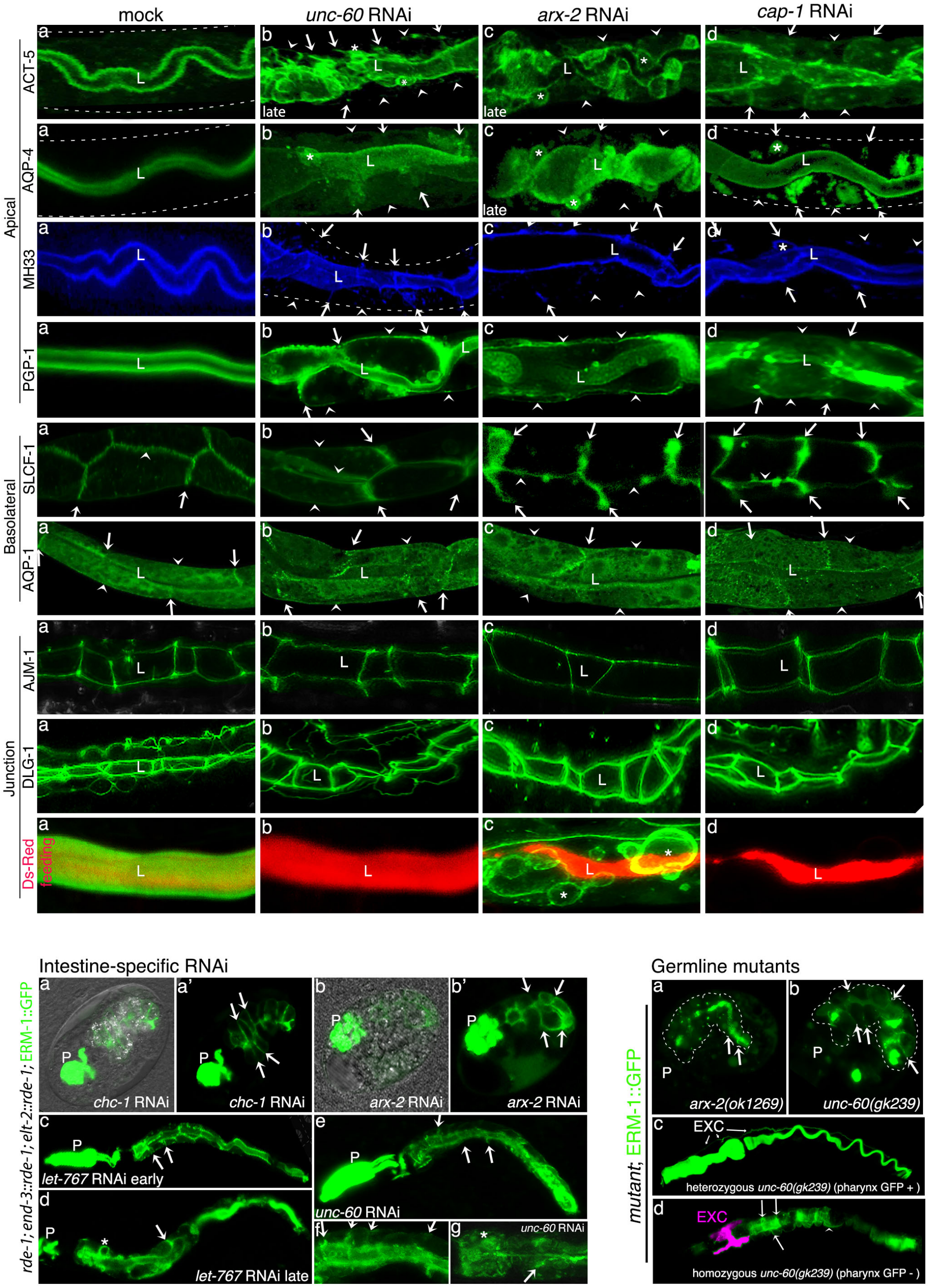

mock
unc-60 RNAi
arx-2 RNAi
cap-1 RNAi
Apical
ACT-5
AQP-4
MH33
PGP-1
Basolateral
SLCF-1
AQP-1
Junction
AJM-1
DLG-1
Ds-Red
feeding
late
Intestine-specific RNAi
rde-1; end-3::rde-1; elt-2::rde-1; ERM-1::GFP
chc-1 RNAi
arx-2 RNAi
let-767 RNAi early
let-767 RNAi late
unc-60 RNAi
Germline mutants
mutant; ERM-1::GFP
arx-2(ok1269)
unc-60(gk239)
EXC
heterozygous unc-60(gk239) (pharynx GFP + )
homozygous unc-60(gk239) (pharynx GFP - )

**Figure S1. UNC-60, ARX-2, and CAP-1 determine the polarized distribution of apical, but not basolateral, membrane components. Intestine-specific depletion of bcAMs and germline mutants (related to Figure 1).**

**Apical membrane panel**: *unc-60* (b), *arx-2* (c), *cap-1* (d) RNAi mislocalizes submembranous (ACT-5/actin; IFB-2/intermediate filament) and integral apical membrane components (AQP-4/aquaporin; PGP-1/P-glycoprotein-related) to basolateral (BL) membranes of expanding larval intestinal cells (a: wild-type localization). Early stages of polarity conversion are shown (otherwise marked as 'late'). L: lumen (apical domain); arrows: lateral portion of BL membrane; arrowheads: basal portion; asterisks: ectopic lumens (ELs). For clarity, intestine is outline by dashed lines in some images.

**Basolateral membrane panel**: *unc-60* (b), *arx-2* (c), *cap-1* (d) RNAi has no effect on the positioning of the integral basolateral membrane components SLCF-1/solute-carrier-family-member and AQP-1/aquaporin (BL membrane structure is affected but not mispositioned). Images for SLCF-1 show the basal portion of the BL membrane (arrowheads; apical domain/lumen is not visible in these focal planes).

**Apical junction panel**: *unc-60* (b), *arx-2* (c), *cap-1* (d) RNAi maintains the peri-lumenal ladder pattern of apical junctions during larval polarity conversion (AJM-1 and DLG-1/discs-large of the junctional DAC complex are shown; [28]. Junction integrity is shown by failure of ingested DsRed bacteria to leak in-between BL membranes or into basolateral ELs (asterisks) or ERM-1+ apical vacuoles (see text).

Confocal sections of 2 pairs of opposing cells of larval intestines are shown. All membrane-/junction markers are GFP fusion, except IFB-2, immuno-stained with MH33/Cy5 (Methods).

**Intestine-specific RNAi.** ARX-2 and UNC-60 function cell-autonomously in the intestine and *arx-2-* and *unc-60* RNAi copy the *chc-1(RNAi)* early-embryonic polarity defect and the *let-767(RNAi)* larval polarity conversion (*chc-1* encodes the clathrin heavy chain and *let-767* a glycosphingolipid-biosynthetic enzyme; both function as vesicle-based apical polarity cues in the *C. elegans* intestine ([5]; see text).
(a-b') Failure of ERM-1 polarization and intercalation in early-embryonic intestine subsequent to intestine-specific RNAi with *chc-1* and *arx-2*. (c-g) Larval polarity conversion subsequent to intestine-specific RNAi with *let-767* and *unc-60*. *rde-1; end-3::rde-1; elt-2::rde-1;* ERM-1::GFP was used with a MYO-2::GFP co-injection marker that marks the pharynx (P) to generate the intestine-specific RNAi strain (Methods).

**Germline mutants.** (a) Embryonic polarity defects in *arx-2(ok1269*). (b-d) Embryonic (b) and larval (d) polarity defects in *unc-60(gk239)*, (c) wild-type (see Fig.S2 for genotypes of balanced mutant strains). Note that ERM-1 is fully displaced from the apical membrane in the anterior intestine in (a). Wild-type and heterozygotes are distinguished by a MYO-2::GFP-positive pharynx (P), therefore, any animal with a GFP-negative pharynx is a homozygous mutant (see Fig.S2 for genotypes of mutant strains).

Confocal images and confocal/Nomarski overlay images (a, b) are shown. Arrows: BL ERM-1 mislocalization. Excretory canal (EXC) in (d) pseudocolored (purple) to distinguish from the intestine.

Figure S2

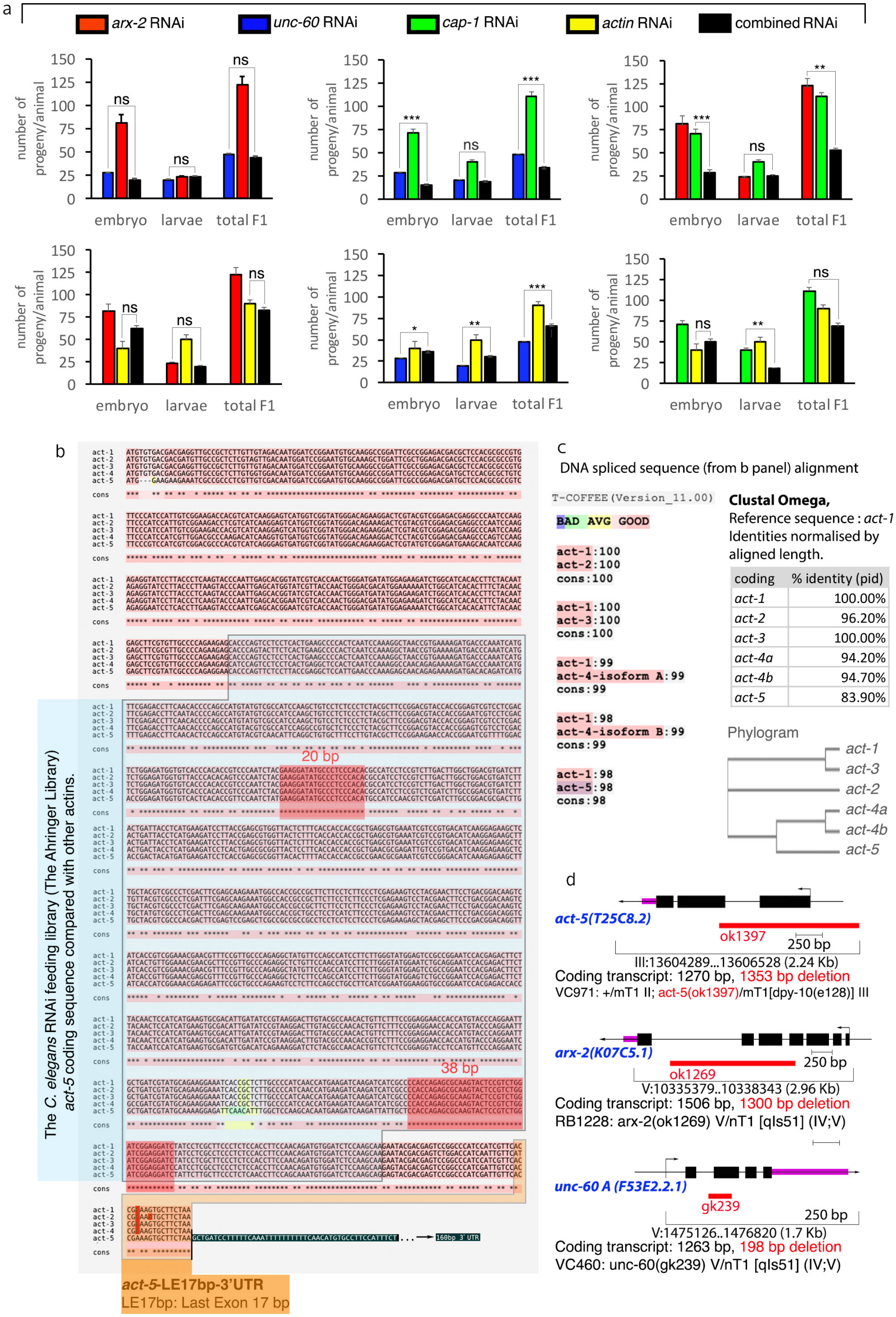
a
arx-2 RNAi
unc-60 RNAi
cap-1 RNAi
actin RNAi
combined RNAi
number of progeny/animal
150
125
100
75
50
25
0
embryo
larvae
total F1
ns
***
**
*
b
20 bp
38 bp
The C. elegans RNAi feeding library (The Ahringer Library) act-5 coding sequence compared with other actins.
act-1
act-2
act-3
act-4
act-5
cons
160bp 3' UTR
act-5-LE17bp-3'UTR
LE17bp: Last Exon 17 bp
c
DNA spliced sequence (from b panel) alignment
T-COFFEE (Version_11.00)
BAD AVG GOOD
act-1:100
act-2:100
cons:100
act-1:100
act-3:100
cons:100
act-1:99
act-4-isoform A:99
cons:99
act-1:98
act-4-isoform B:99
cons:99
act-1:98
act-5:98
cons:98
Clustal Omega,
Reference sequence : act-1
Identities normalised by aligned length.
coding
% identity (pid)
act-1 100.00%
act-2 96.20%
act-3 100.00%
act-4a 94.20%
act-4b 94.70%
act-5 83.90%
Phylogram
act-1
act-3
act-2
act-4a
act-4b
act-5
d
act-5(T25C8.2)
ok1397
250 bp
III:13604289...13606528 (2.24 Kb)
Coding transcript: 1270 bp, 1353 bp deletion
VC971: +/mT1 II; act-5(ok1397)/mT1[dpy-10(e128)] III
arx-2(K07C5.1)
ok1269
250 bp
V:10335379..10338343 (2.96 Kb)
Coding transcript: 1506 bp, 1300 bp deletion
RB1228: arx-2(ok1269) V/nT1 [qIs51] (IV;V)
unc-60 A (F53E2.2.1)
gk239
250 bp
V:1475126..1476820 (1.7 Kb)
Coding transcript: 1263 bp, 198 bp deletion
VC460: unc-60(gk239) V/nT1 [qIs51] (IV;V)

**Figure S2. Genetic interactions among branched-chain actin modulators (bcAMs) and between bcAMs and actin in lethality and sterility. Comparison of exonic sequences of *act-1, -2, -3, -4,* and -5 and areas targeted by a commonly used *act-5* RNAi clone. Germline mutant strains (related to Figure 2).**

(a) Total number of progeny and arrest stages (embryonic versus larval) are shown. RNAi with each bcAM and actin cause lethality and sterility (# of N2 wild-type progeny is approx. 220 at 20$^0$C, [3 29]. At least 3 replicas were analyzed. *t-test* was used to compare the significant value of each pair. Compare to Figure 2 (interactions in lethality do not simply reflect those in polarity (see below* for details).

(b) The targeted area in *act-1-5* of the broadly used *C. elegans* RNAi feeding clone *act-5* (The Ahringer Library; [10] is indicated by light-blue overlay over spliced sequence alignment (small areas of relevant discrepancy: yellow; see (c) for color coding). The targeted area in *act-1-5* of the *act-5-LE17bp-3'UTR* RNAi clone (see text) is indicated by orange overlay (discrepant bases highlighted in bright red). Dark pink-red overlay shows conservation of more than 18bp, considered necessary for effective RNAi in *C. elegans*. Note that the *act-5-LE17bp-3'UTR* dsRNA targets an additional stretch of 17 base pairs of exonic sequence of which 14-16 match *act-1-4*. Although below the cutoff of 18 base pairs thought to be required for full RNAi effect, *actin-* and *act-5-LE17bp-3'UTR-,* but not *act-5-3'UTR* RNAi, reduce expression of ACT-1- and ACT-2::GFP (see Fig.7 and not shown).

(c) Comparison of the almost identical *act-1-5* exonic sequence scores. The color coding on top of the panel shows the level of conservation, as defined by T-COFFEE [30]; compare to Clustal Omega. DNA sequence maximum-likelihood phylogenies shown beneath.

(d) Genomic organization of *act-5*, *arx-2* and *unc-60* and genotypes of germline mutant strains. All genes have exonic deletions and are predicted or demonstrated to be genetic nulls. *act-5(ok1397)* also deletes the promotor; *unc-60(gk239)* removes the UNC60A isoform that is expressed in embryonic and non-muscle tissues, including the intestine; [31]. Note that all genes are maternal-effect embryonic/larval lethal, and the required balancers still provide maternal product to the homozygous mutant progeny.

*Note that *unc-60* dependent lethality was not significantly suppressed by *arx-2-*, but mildly by *cap-1* RNAi, whereas *cap-1(RNAi)* lethality was enhanced by *arx-2* RNAi. More similar to bcAMs' inability to enhance actin's polarity defects, *arx-2* and *cap-1* RNAi also failed to enhance the actin-dependent lethality. Actin loss, however, mildly suppressed the lethality of *unc-60* RNAi, suggesting that imbalances in G- and F-actin stoichiometries themselves (expected to occur by perturbing *unc-60* mediated filament disassembly) induce toxicity (compare cartoons of bcAM interaction in Fig.2).

**Anterior-posterior apical membrane extension during intestinal development**

a

| | Length | SEM |
|---|---|---|
| Bean | 21.62 | 0.3 |
| L1 | 80.26 | 2.822 |
| L2/3 | 243.6 | 8.303 |
| L4 | 684.6 | 15.49 |
| Adult | 735.1 | 32.31 |

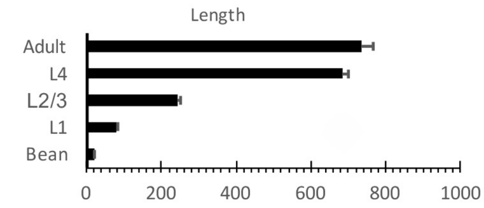


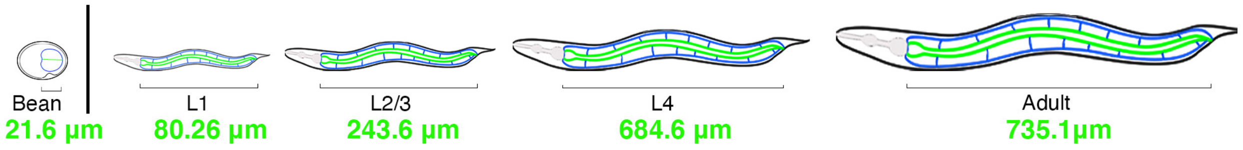


b

| | Length | SEM |
|---|---|---|
| Bean | 21.62 | 0.3 |
| Comma | 24.71 | 0.288 |
| 1.5 fold | 35.92 | 0.27 |
| 2-fold | 54.1 | 0.608 |
| 3-fold | 67.05 | 0.894 |

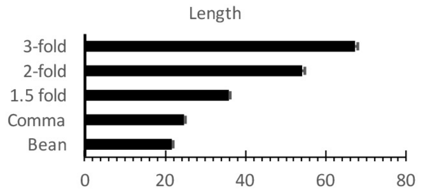


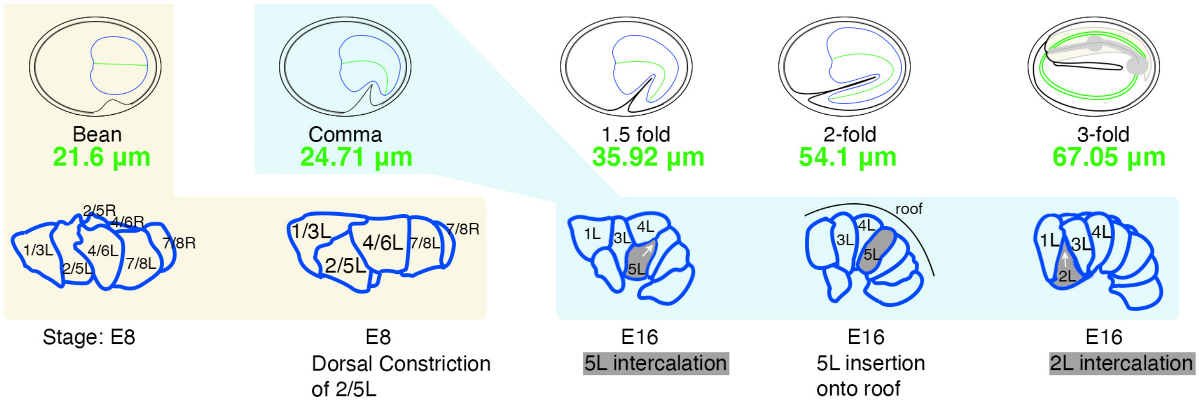


**Figure S3. Embryonic and larval intestinal development and net extension of anterior-posterior length of the intestinal apical (lumenal) membrane.**

(a) Increase in apical membrane length from 80.26 µm to 735.1 µm from L1-larva to adult, corresponding to an increase from 8.9 to 81.7 µm per cell (9 cells per row: 20 cells form 9 INT rings consisting of 2 cells each, 4 cells in INT 1). Schematics show fully formed intestine that extends through 4 larval stages to the adult (only minimal further extension in adults). Here and below: apical membrane: green, basolateral membrane: blue; anterior left, posterior right; dorsal up, ventral down; R/right left, L/left right.

(b) Increase in apical membrane length during embryogenesis from 21.6 µm (7 cells per row; bean stage) to 67 µm (9 cells in one row; 3fold stage), corresponding to an increase from 3 to 7.4 µm per cell (taking intercalation into account = the addition of 2 cells per row; see below). Schematics show intestinal morphogenesis from bean to 3-fold embryo, with early-embryonic pre-intercalation (E8 stage; light yellow shadow) and intercalating intestines (E16 stage; light blue shadow) shown below (adapted from [32]. The left row of cells is shown (L). Intercalating cells are shaded in dark grey. Orientation as above, dorsal side (roof) is indicated.
Note that these measurements do not take into account the large amount of membrane required for the formation of microvilli at the apical intestinal membrane.

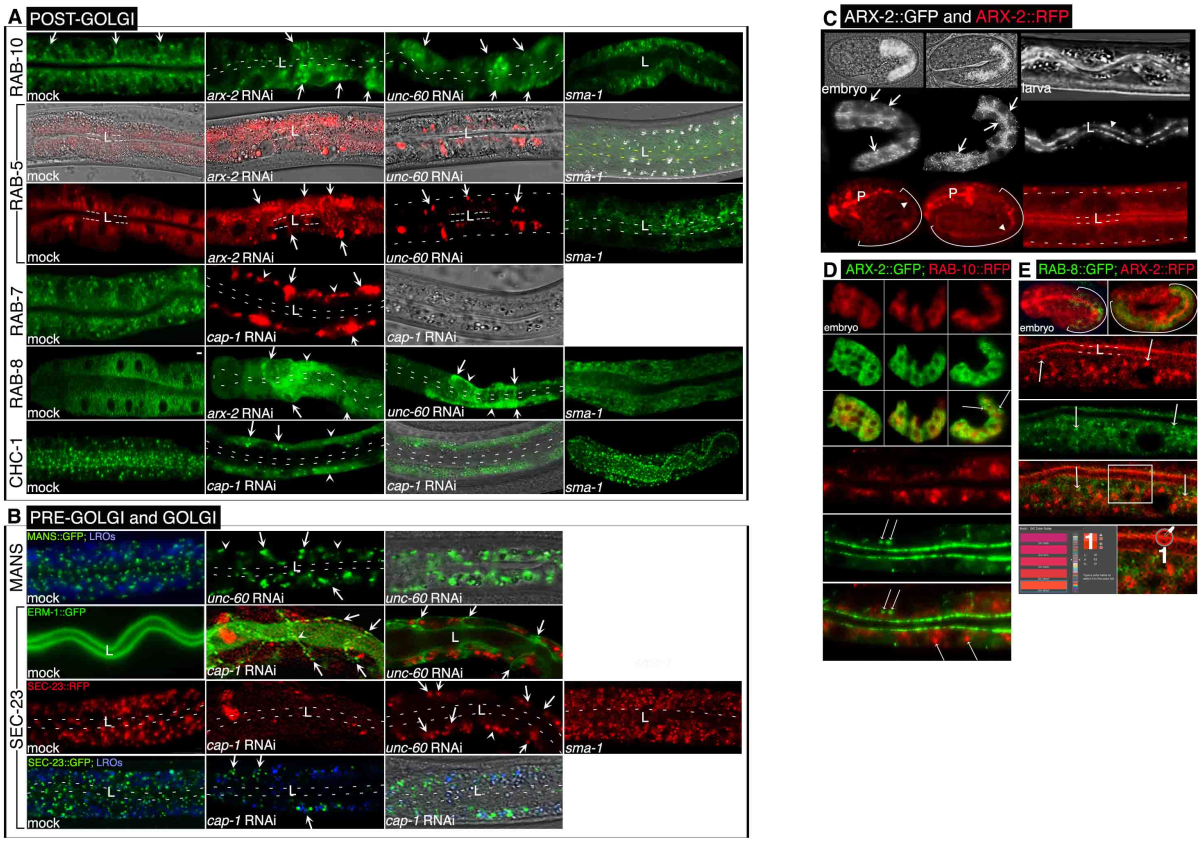


**Figure S4. The apical targeting of pre- and post-Golgi vesicles during *de novo* polarized membrane biogenesis depends on UNC-60, ARX-2, CAP-1, and actin (related to Figure 6).**

(A-B) Post-Golgi, Golgi-, and pre-Golgi endomembranes/vesicles fail to reach the apical domain (lumen/L, arrowhead, indicated by dashed lines where difficult to distinguish from background) and are misdirected to basolateral (arrows) and basal (arrowheads) membranes of growing *unc-60-, arx-2-, cap-1 and actin-,* but not control *sma-1(RNAi)*, late-embryonic/early-larval intestinal cells (see text and compare to Fig.6). Corresponding Nomarski or Nomarski overlay images reveal morphologically intact intestines. ERM-1::GFP basolateral displacement in ERM-1::GFP SEC-23::RFP double labeled intestine reveals concomitant membrane polarity change (basolateral to apical).

(C) ARX-2 expression: intestinal ARX-2::GFP expressed from a transgene recapitulates the embryonic (left) and larval (right) expression pattern of ARX-2::RFP expressed from its germline locus, but better resolves ARX-2 speckles in the intestine (compare Fig.4). Basolateral membranes indicated by arrows, embryonic intestine bracketed, P: pharynx, L/arrowhead and double lines: lumen, dashed line: basal intestinal membrane.

(D) ARX-2::GFP speckles fail to colocalize with RAB-10::RFP+ vesicles (thin arrows), consistent with ARX-2::RFP GFP::RAB-10 double labeling (Fig.6).

(E) ARX-2::RFP speckles also fail to colocalize with RAB-8::GFP+ vesicles (thin arrows). Lower panel shows magnified merged portion of double labeled intestine, measured by the Eyedropper tool (Adobe-22.4.3 DIC color guide; compare Fig.6). ARX-2::RFP brightness increased to attempt to resolve speckles (compare Fig.4).

Confocal and confocal/Nomarski overlay images of full embryonic and partial late-embryonic/early-larval intestines (2 pairs of opposing cells) are shown. See Methods for fluorescent fusion proteins.

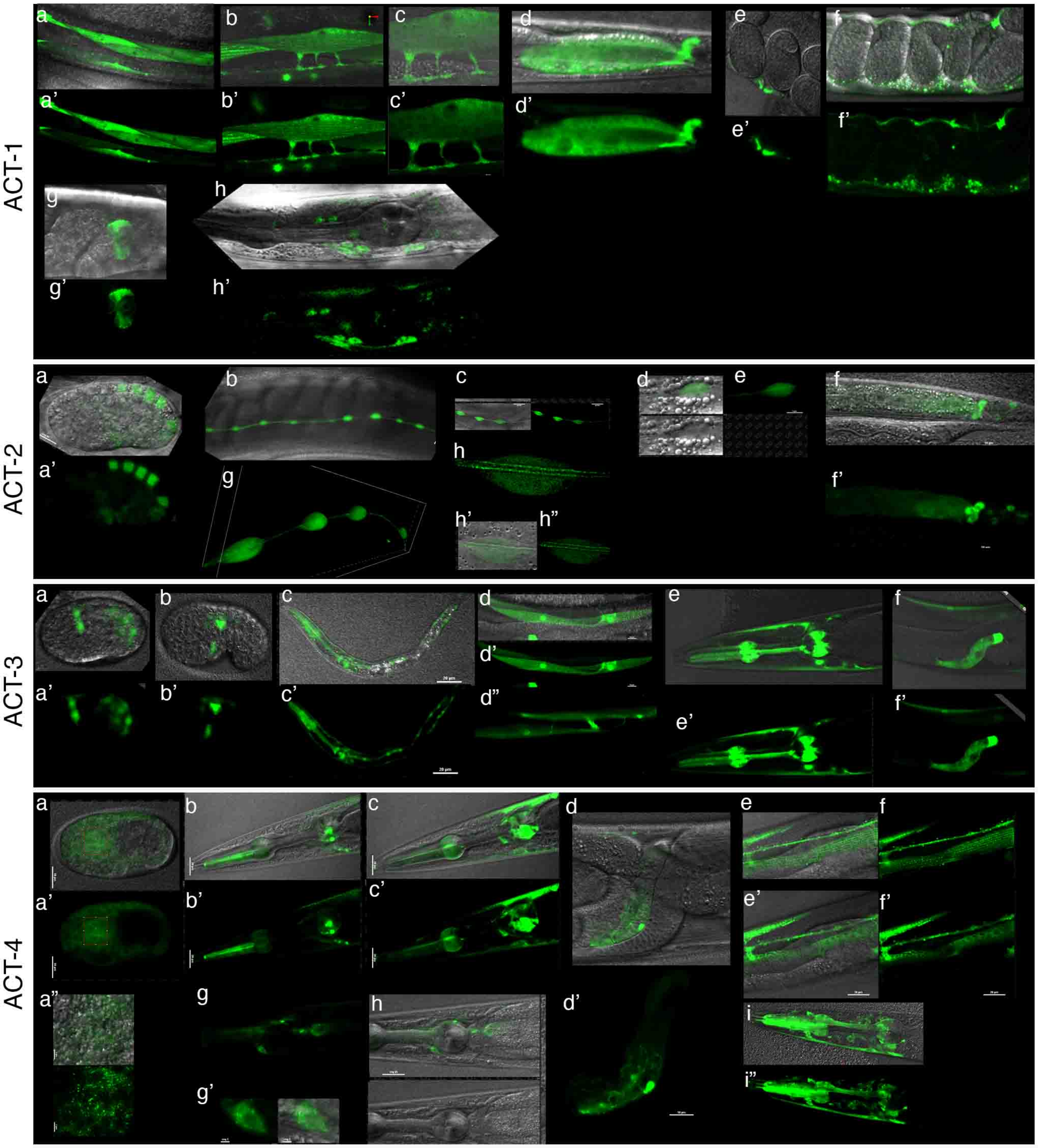
ACT-1
ACT-2
ACT-3
ACT-4

**Figure S5. Non-intestinal expression profile of ACT-1, ACT-2, ACT-3, and ACT-4 (related to Figure 7).**

**ACT-1::GFP:** Expression in: (a,a') body wall muscles; (b, b', c, c') muscle extensions (muscle arms) that form *en passant* neuromuscular junctions (note ACT-1 puncta); (d,d') adult intestine and gland cells; (e, e') vulval muscles; (f, f') intra-uterine speckles; (g, g') coelomocyte; (h, h') head neurons.

**ACT-2::GFP:** Expression in: (a, a') embryonic hypodermis; (b, c) seam cells (magnified in d, e, 3D image in g; with alae in h-h"); (f, f') adult intestine and gland cells.

**ACT-3::GFP:** Expression in: (a, a', a", b, b') embryonic muscle cells; (c, c') pharyngeal, body wall muscles and other tissues (full view of an L2 larva); (d-d") muscles and muscle extensions (muscle arms); (e, e') pharyngeal and body wall muscles; (f, f') spermatheca.

**ACT-4::GFP:** (a, a') Expression in most embryonic tissues except the intestine (note speckled pattern in a"); (b, b', c, c') pharyngeal, head neurons (magnified in g, g') and body wall muscles; (d, d') spermatheca; (e, e', f, f') ventral nerve cord, muscle, and muscle arms; (h) head neurons; (i, i') body wall muscles, pharynx, and head neurons.

Confocal and confocal/Nomarski overlay images are shown. See Methods for fluorescent fusion proteins.

Only partial expression profile is shown (full profile in Jafari et al., in preparation).

**a**

LAST EXON END, stop codon (TAA), off-target 3'UTR.
different colores cloned RNAi "on target" 3'UTR.

*act-1* 3'UTR sequence, on target 596 bp

TCACCGCAAGTGCTTCTAAatgcacaacttcgtcaacttgcacaaacaagttcaaaaccatcgtcaccaccagctttctatttcttgtttgcatatttttgtgcaacaaggaaacatcatgcatatttcccaaaa aaataaaaaaccgccccatttttgtccttattgtcctttcactcatttgcgggtgtgaataaaaatttaaaaattgatatttctaaattgtctatcagcaccattaacagattcttgaagcactcgccgccaaaaaatgttactgt ttgagaaatacgttgaagacagttgcgacatgagagatatttgagtgggcgcagagcgattccgagaatcgcaacgttcgtgaagatactgctcgtctctcgagaaagaaagcacacattccatagtttcttcttattcc cacggcggcgaaagcatctgtttacacaacatgcggaggcaaaaacgacgagaagcgtcgaactgataggaagaaaacagaaaatctttgaacaggaagggaagtgtaggcggttgtatgataacttccgtac ggataatagttttaaatgtaaacaatgttaattaagaacacaaacttctccattgattcaa

*act-2* 3'UTR sequence, off target, on target 331 bp

TTCATCGCAAATGCTTCTAAacgttttaacaatttatgtaatattcacctgaacttttggaaatgtccagataacgtcatctgccttacggattttcacaccccgattttaattataaatctgcttcgtgtaagcatcgt tttaaaatttgtcggtttattttaataatatttctaaacgtgaaatcgtatatatacttttagtttagtggtccaatgtcatcattggtaaacttcatcgtttaacgtgacctccaagtccagatcgcggttgttgataagcatttctggag tcaacttcttatcagttcggtgtcgattgcgttcactgttcctgttgattgaaatttcccgcgttacgataatctcgtctctccctcttcgccagttgctctttcgggccgaaattttctttatctaactgtttc

*act-3* 3'UTR sequence, off target, on target 196 bp

CACCGCAAGTGCTTCTAAgctcttcgccttaccattttctcttttcttttctttttttctatacattttttttccaattgtcaccaaatcgttatggtccctttttgggg caataacaaattctatcatcatttcacgatcatgagaccattcaaaa agaccatcaagattttatttccgatcctgccacataagaagaccccactacaatttgctctgttctttgttttgtgcatatttgtttttttaattgcttcttgctatacacaaaatttacaaaataaaacgttccgggctcattttctgg

*act-4* 3'UTR sequence, on target 332 bp

GTCCACCGCAAGTGCTTCTAAatttttgccccttccacccacaacgctcaaaagaagttcgtattatcccgcaacccatccagaccatccttttacatttttttatttctcttttttttttcttcaccctctccgtgtaca ttgctagtaatagttccaacttctctagccgactgcattatgcttcccattctcacccaaatcttcattggtaacttctcttgtatttgttcaatactattccggtagtaatctaggcatgtataactaggttctctgtgtatctttggtttga agttgaggtttaatgcaatcacttttgttgttgtaccctattactgccgatgacc

*act-5* 3'UTR sequence, on target 148 bp

CCGAAAGTGCTTCTAAgctgatcctttttcaaattttttttttcaacatgtgccttccatttctaggcgaaacacgattttacttgacacaaattccatgagatacttcattttctttatatttatggtttctt aacatgaataaacttgttaggaccacttcg

| | |
|---|---|
| *act-1* | *V:11081052..11082415* |
| *act-2* | *V:11078154..11076777* |
| *act-3* | *V:11074498..1071690* |
| *act-4 a* | *X:4960621..4964413* |
| *act-4 b* | *X:4962857..4963982* |
| *act-5* | *III:13606066..13604412* |

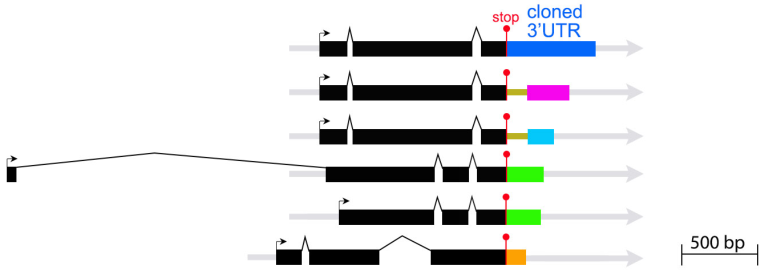


**b**

*act-1(tm6888)*

*act-2(ok1229)*

*act-3(tm6924)*

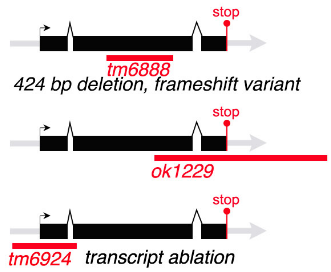


**c**

**Promoter comparison.**
**2 KB analysed**

BAD AVG GOOD

T-COFFEE software controls:
(-) Control, two unrelated genes:
*act-1* vs *mab-9*

```
control-mab-9   :  22
```

(+) Control: two identical genes
*act-1* vs *act-1*

```
act-1P-mock   : 100
```

Comparing *act-1*, *act-2*, *act-3*, *act-4a*, *act-4b*, *act-5* promoters

```
act-1    :  16
act-2    :   9
act-3    :  10
act-4A   :   8
act-4B   :  10
act-5    :   9
```

**Intron comparison.**

First intron *act-1 vs act-2 vs act-3*

```
act-1  :  96  GTAAATAATTAATACATTCGAT-GATTAAATTTATGCGTACTATTTCAG
act-2  :  94  GTAAATTTTCAAAAAATTTGACCGATTGGAAATA---GTTGTTTTT-AG
act-3  :  97  GTAAATAATTAATACATTCGAT-GATTAAATTTATGCGTACTATTTCAG
              ******  * ** * *** **  ****  *  **   **  * *** **
```

First intron *act-1 vs act-4a vs act-5*

```
act-1
act-4a  :  23

act-1
act-5   :  22
```

Last intron comparison:

```
act-1  :  82  GTAAGTTTTC--TATTTTTGTTTCAGTTAACAA-ATTAAATTGATTTTTTTTT-CAG
act-2  :  87  GTAAGTTTTC--TATTTTTGTTTTAGTTAACAA-ATTAATTTGATTTTTTTTT-CAG
act-3  :  90  GTAAGTTTTC--TATTTTTGTTTCAGTTAACAA-ATTAATTTGATTTTTTTTTCAG
act-4  :  77  GTAAGTTTAACTTAAATGGATTTCAAGAACTAATATTTACTTTC-TATTTTTTCAG
act-5  :  79  GTAAGCTTCA--CATTT----CTCAGTACATGAGATTATAATTCTTGTATTTTCAG
              ***** **      *  *      * *        * ***    *   * * *** ***
```

**3'UTR comparison.**
**475 bp analysed.**

```
act-1-3utr   :  20
act-2-3utr   :  13
act-3-3utr   :  12
act-4a-3utr  :  22
act-4b-3utr  :  24
act-5-3utr   :   8
```

**d**

**Single-cell transcriptional profiling; Intestinal cells.**

| | |
|---|---|
| *act-1* | 34.31 |
| *act-2* | 6.89 |
| *act-3* | 51.42 |
| *act-4* | 86.69 |
| *act-5* | 1210.85 |

Values listed are transcripts per million.

**Figure S6. Sequences targeted by RNAi vectors generated for the 3'UTR RNAi experiments (see Methods for rational of approach). *act-1-3* germline mutants. Comparative analysis of intron/exon structure and of non-coding regions of *act-1-5* (promoters, introns, 3'UTRs). RNAseq analysis of *act-1-5* in the intestine (see Fig.S2 for exonic *act-1-5* sequences, Fig.S7 for sequence alignment of ACT-1-5 proteins).**

(a) Each 3'UTR targeted in this study is highlighted in a different color. Please note that any possible off-target areas of *act-2* and *act-3* (olive) were excluded when generating the RNAi vectors (Methods). Using these dsRNAs, specifically designed to avoid off-target effects, neither *act-1-* or *act-4-3'UTR* RNAi, nor *act-1+2-3'UTR-* and *act-2+3-3'UTR* double RNAi, nor *act-2(ok1229) act-1+3-3'UTR* double mutant/RNAi induced lethality (compare to [8]).

The lower panels show intron/exon structures of *act-1-5*, relative to the 3'UTRs. *act-1, 2* and *3* are located on a cluster on chromosome V, the other actins on other chromosomes. Only 1 alternatively placed isoform is predicted (*act-4*).

(b) Germline deletions of *act-1-3*. Strains are homozygous viable.

(c) T-COFFEE alignment of promoters, introns, and 3'UTR of the 5 actin isoforms. (-) and (+) controls are shown.

(d) Transcriptional profiling of actin isoforms in intestinal cells by RNAseq [33].

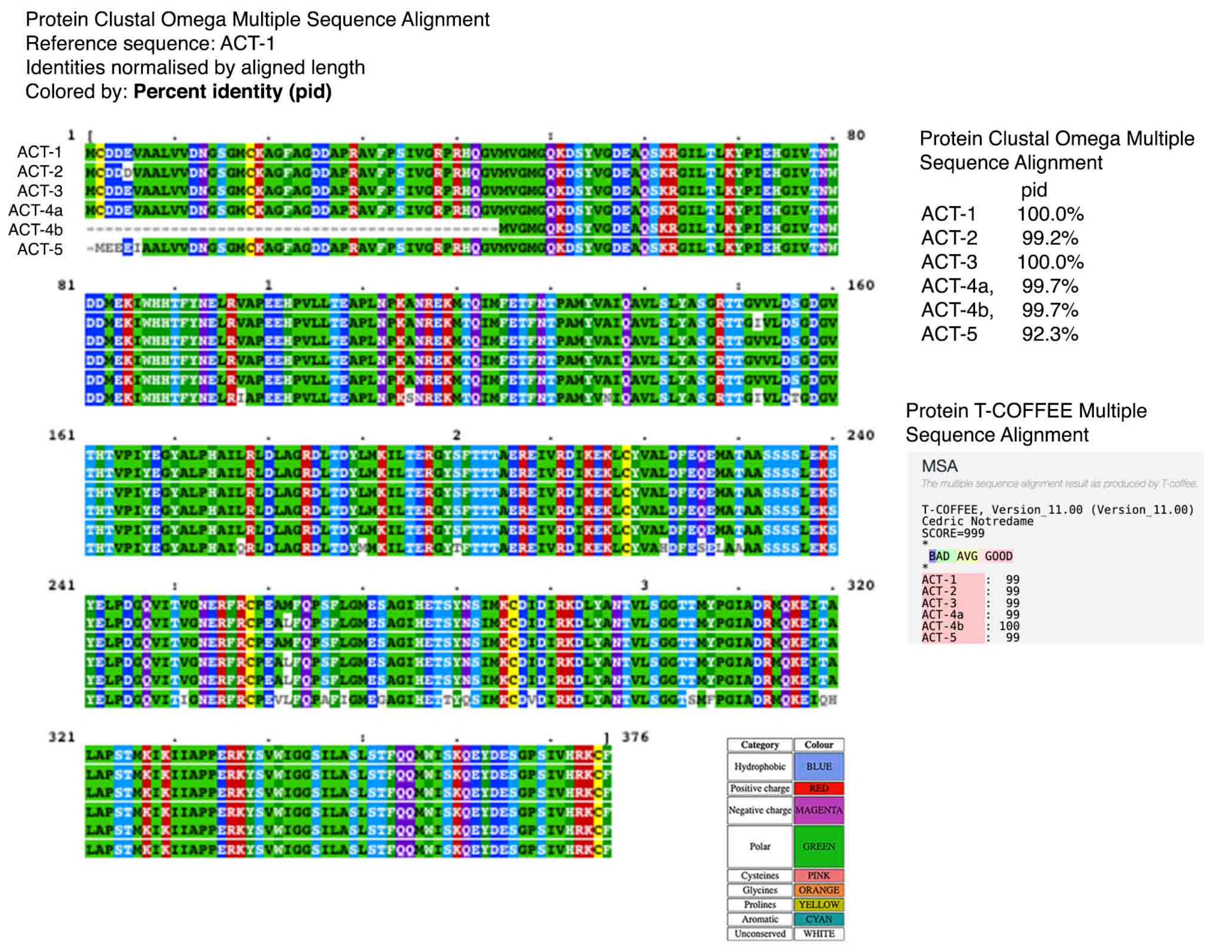


**Figure S7. Sequence alignment of ACTIN proteins**.

ACT-1 and ACT-3 are fully identical, distinguished by 2 amino acid changes from ACT-2; by 1 amino acid change from ACT-4 (each of 2 isoforms); by 24 amino acid changes from ACT-5. Comparison of Clustal Omega and T-COFFEE alignments is shown.

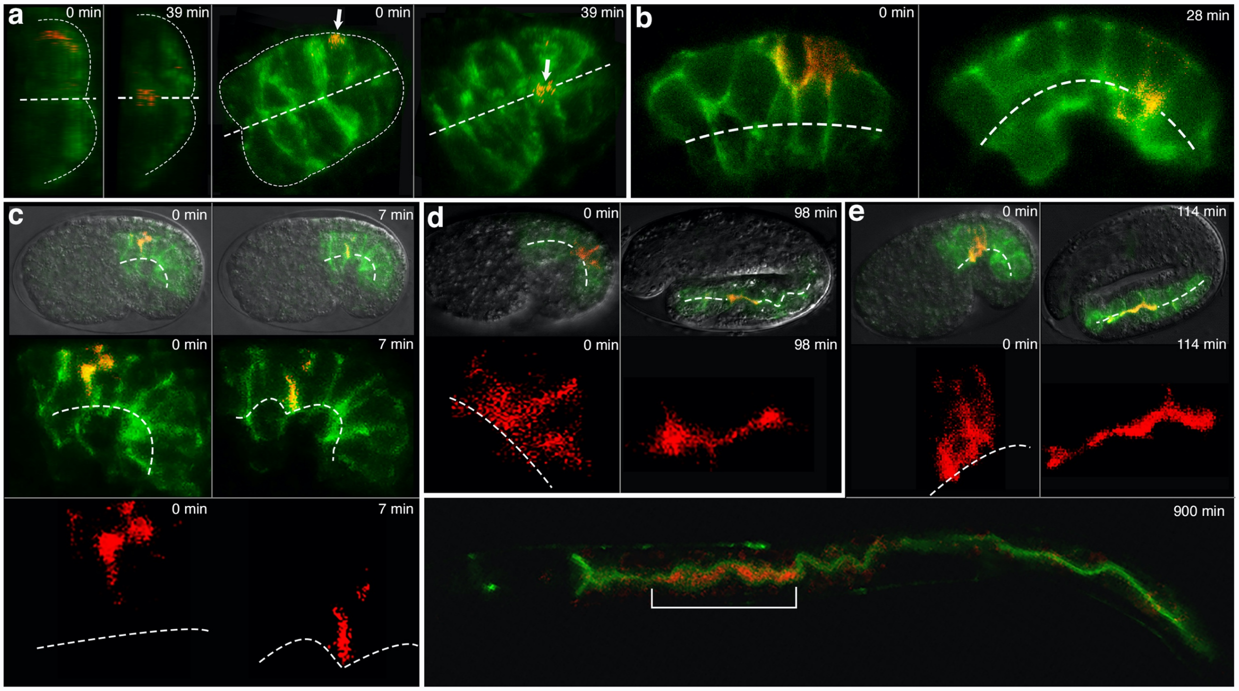


**Figure S8. Tracking the transcytotic F-actin shift from the basolateral to the apical domain in single intestinal cells with time lapse imaging of photoconverted LifeAct::Dendra2 (related to Figure 9).**

See Fig.9 and Methods for photomodulation procedures. Note that LifeAct::Dendra2 is converted from green to red (allowing for concomitant tracing of both converted [red] and non-converted [green] F-actin, making this color scheme different than that of PA-GFP (Fig.9).

(a) In the first two panels, ventral is to the left and dorsal is to the right. Photoconversion of basal portion of basolateral membrane of a single cell of the pre-intercalation intestine (early bean stage) during polarity establishment. In the third and fourth panels, a dorsal view; anterior is to the left and posterior to the right (compare Movie 1). Note that only converted LifAct::Dendra2 is red and thus translocates from basolateral (red, 0 min) to the midline (dashed line: future lumen; red, 39 min). This embryo is rotated in Movie 1.

(b) Photoconversion of basolateral membranes of two cells at later stage of development (late intercalation; comma stage), revealing ongoing F-actin translocation to the apical domain (dashed line). Lateral view: anterior is to the left and posterior is to the right.

(c) Tracking photoconverted LifeAct from basolateral membrane of one cell (late bean stage) at shorter time interval reveals basal to lateral F-actin translocation. This embryo is rotated in Movie 2.

(d) Photoconversion of lateral membrane of one cell in post-intercalation intestine (1.5fold embryo) reveals lateral to apical F-actin translocation.

(e) Tracking photoconverted LifeAct from the latero-apical angle of E2 and E3 (converted at late stage of intercalation) through longer time interval reveals full translocation of LifeAct into the apical domain in hatched larva (900 min).

**Movie 1. A 3D rotating view of the embryo from Figure S8, a.**

**Movie 2. A 3D rotating view of the embryo from Figure S8, c.**

### Supplementary Results

**ACT-5 is not the only actin in the *C. elegans* intestine**

To molecularly characterize an actin structure able to constrain anterograde trafficking during membrane polarization, we explored if ACT-5 was sufficient to execute actin's polarity function (a question raised by the absence of embryonic polarity defects and lethality in *act-5(ok1397)* mutants and by the possibility that *'act-5'/actin* RNAi might have targeted other, highly similar actins; Figs.2, 2Sb-c). Indeed, the LifeAct imaging analysis had suggested a broader subcellular F-actin distribution than that of ACT-5 (Fig.4), so far considered restricted to the apical membrane and its microvilli based on rescuing ACT-5::GFP- and ACT-5::mCherry fusions (detecting both F- and G-actin pools [34]; also based on the epifluorescent and confocal microscopic analysis of the high affinity F-actin chemical stain phalloidin and pan-F-actin). Moreover, RNAseq identifies the expression of all other *C. elegans* actins (ACT-1-4) in the intestine, albeit at low levels (Fig.S6d). We therefore first asked if other actin isoforms might be present in the intestine and generated transgenic animals with ACT-1-4::GFP fusions to re-assess the localization of these isoforms by high resolution confocal microscopy (Methods/Fig.7 legend for rationale for transgenesis).

As expected, ACT-1-4::GFP were all strongly expressed outside the intestine, e.g., in the hypodermis during early embryonic development (ACT-2) and in muscle and muscle attachment structures throughout life (ACT-1 and 3; Fig.S5). However, ACT-1-4::GFP were also expressed in the intestine, albeit weakly (Fig.7; note that confocal image acquisition is adjusted, and brightness increased for ACT-1-4, but not for the strongly expressed ACT-5). A comparative subcellular developmental expression analysis of all actins revealed that ACT-5::GFP is also expressed in the intestinal cytoplasm, especially in the pre-intercalation intestine, and fully recapitulates the basolateral-to-apical shift of LifeAct::GFP during polarity establishment in the early embryo (Fig.4). Moreover, ACT-1-3::GFP, although mainly located in the cytoplasm, are also enriched at the cortex of early-embryonic intestinal cells and, during polarization, synchronously shift position from the basolateral to the apical membrane (especially ACT-2 which is also expressed in the excretory canal; not shown). ACT-4::GFP could only be detected in the posterior adult intestine. We conclude that: (1) ACT-5, although the prominent actin, is not the only actin in the *C. elegans* intestine; (2) ACT-5, although chiefly located at the apical membrane, is also located in the cytoplasm and at all sides of non-polarized membranes in the early-embryonic intestine; (3) cortex associated ACT-5 undergoes a basolateral-to-apical polarity shift during intestinal polarization that is copied by ACT-1-3. LifeAct captures the subcellular location of all 5 actins, with strong cytoplasmic expression and an additional perinuclear, punctate (early embryo) and peri-vesicular location (larva and adult; see Fig.7 legend for further details; see below and Fig.8 for a high magnification analysis of intestinal actin). The expression of all 5 actin isoforms in the intestine and the synchronous basolateral-to-apical positional shift of ACT-1, -2, -3 and -5 during membrane polarity establishment raised the possibility that several actin isoforms might contribute to an actin structure proposed to direct anterograde vesicle trajectories towards the expanding apical domain during membrane polarization.

**ACT-5 interacts with other actin isoforms in apicobasal membrane polarity**

To test the hypothesis that actin isoforms other than ACT-5 were required for actin's polarity function, we first examined if polarity conversion could be induced by interference with ACT-5 alone. To target maternal product, avoid full depletion (to preserve cellular morphogenesis), and prevent targeting of the highly similar *act-1-4* transcripts (Fig.2S), we tested *act-5* specific 3'UTR RNAi. *act-5-3'UTR* RNAi only induced mild lumenogenesis but no polarity defects (we failed to reproduce the previously reported L1 lethality [Waddle] with an *act-5-3'UTR* dsRNA designed to minimize off target effects; Fig.S6a and Methods). An "allelic series" of: (1) *act-5 3'UTR* RNAi (Fig.S6a), (2) *act-5* (*ok1397;* presence of maternal product; Fig.S2d), (3) *act-5 LE17bp-3'UTR* RNAi (possibly targeting other actins; Fig.2Sb) (4) *actin* (aka *act-5* RNAi, commonly used with the intent to target *act-5* but in fact targeting all actins; Fig.2Sb) displayed increasingly severe lumenogenesis and apical membrane polarization defects. However, apicobasal polarity defects (basolateral displacement of the apical domain) were only present in (3) and (4), conditions distinguished by their ability to target other actins (Tab.S1, Figs.7, S2b). Finally, ACT-5 overexpression produced L1 lethality and a strong lumenogenesis, but no polarity, phenotype. We concluded that *act-5* was likely contributing to actin's polarity function but might not be sufficient for it.

To ask if *act-5* was even necessary for actin's function in polarity, we moderately increased the strength of RNAi conditions (Methods; see Fig.S6 legend for rationale of choosing mild rather than strong loss- or gain-of-function approaches for the genetic analysis of actin isoforms). An RNAi sensitive *rrf-3* background increased all phenotypic effects of *act-5 3'UTR-*, *act-5-LE17bp-3'UTR-*, and *actin* RNAi, with full penetrance of the apicobasal polarity phenotype in the latter two. It also induced polarity conversion and L1 lethality in an *act-5 3'UTR(RNAi)* background (Fig.7, Tab.S1). Thus, ACT-5 is indeed required for polarity; maternal and zygotic products are necessary for this function; and additional actin isoforms may be needed for full effect. The results also validate the 3'UTR RNAi approach and demonstrate that its effect is mild and thus suited for the analysis of actin's specific function in membrane polarity (see above).

ACT-5 is the most divergent of the 5 highly similar *C. elegans* actins, with ACT-1-4 presumed to function redundantly in muscle and other extra-intestinal tissues (predictions are based on mutational, not loss-of-function, analyses [35-40]; lack of redundancy was only demonstrated for *act-5* and *act-1* [34]; Figs.S2, S6, S7 for *act-1-5* comparative analysis of: genomic organization; spliced sequences; non-coding sequences; and encoded proteins; legends for discussion). To assess the requirement of the other highly similar isoforms for intestinal polarity, we first examined germline deletions of ACT-1, 2 and 3, the actin isoforms that copied ACT-5's basolateral-to-apical polarity shift during intestinal polarization (Figs 7, S6b). *act-1(tm6888)*, *act-2(ok1229)* and *act-3(tm6924)* were superficially wild-type, consistent with expected redundancies. We did, however, find mild apical domain polarization, albeit no positioning (polarity), defects in these mutants when supplemented with the ERM-1::GFP transgene (distinct patterns of cytoplasmic ERM-1 displacement; not shown). The lack of polarity defects in these *act-1-3* presumed null mutants (Fig.S6b) suggested that, if intestinal actins other than ACT-5 were required for polarity, these actins either require ACT-5 to exert their effect, act redundantly, or, if not redundant, require more than one of these actins for full function.

To distinguish between these possibilities, we targeted the different actin isoforms, either by themselves or in combination, using mild depletion via 3'UTR RNAi (*act-1-5* transcripts are closely similar, while 3'UTRs fully diverge; Figs.S2, S6a, c; Methods; Fig.S6a for dsRNA design; Fig.S6 legend for rationale of trialing this 3'UTR approach). Targeting the 3'UTRs of *act-1-4*, with or without *rrf-3*, did not induce intestinal polarity defects, nor did it enhance the polarity defect of *act-5-LE17bp-3'UTR* RNAi (Fig.7, Tab.S1). The assessment of all double and triple combinations of *act-1-4-3'UTR* RNAi also failed to generate intestinal polarity defects or other gross phenotypes (Fig.7; Tab.S1; Fig.S6 legend for details). In contrast, quadruple *act-1+act-2+act-3+act-4-3'UTR* RNAi induced embryonic and L1 arrest with pronounced elongation and body morphology defects (Fig.8), a widened intestinal lumen, and complete failure of intracellular apical membrane (lumen) expansion in the excretory canal; yet did not induce intestinal polarity defects (not shown). Quadruple *act-1-4-3'UTR* RNAi did, however, induce polarity conversion in an *act-5-3'UTR(RNAi)* background and enhanced it in *act-5-LE17bp-3'UTR(RNAi)* intestines (Fig.7, TabS.1). These results: (1) demonstrate that *act-1-4* interact with ACT-5 in intestinal polarity; (2) suggest that *act-1-4* require *act-5* for this function; (3) reveal that *act-1-4* are, at least in part, non-redundantly required both in and outside the intestine (total actin removal remains constant in single and multiple knockdowns); (4) validate the effectiveness of a multiple (up to quintuple) RNAi approach.

To identify the actin isoform fingerprint for polarity, we next assessed if the combined additional loss of 2 isoforms could induce or enhance the *act-5*-dependent polarity defect. *act-1+act-2+act-5-* as well as *act-3+act-4+act-5-3'UTR* RNAi, but no other combination, increased the *act-5-3'UTR(RNAi)* lumenogenesis defect and induced basolateral displacement of ERM-1; however, these and all other triple 3'UTR combinations with *act-5,* except *act-1+act-3,* were able to increase the *act-5-3'UTR(RNAi)* polarity defect in an *rrf-3* background (Fig.7). We conclude that any combination of at least 2 different actin isoforms (except for the fully identical ACT-1 and ACT-3) are non-redundantly required to cooperate with ACT-5 to exert actin's full polarity function in the *C. elegans* intestine. These findings supported the hypothesis that several actin isoforms might be required to generate an actin structure proposed to asymmetrically direct vesicles to the membrane at the time of its polarized expansion.

#### Supplementary Discussion

A variety of actin assemblies have been implicated in the directional movement of organelles, including that of polarized vesicles. However, the here described bcAM powered actin assembly not only propels vesicles into a given (e.g., apical) direction, but confers (apical) directionality to anterograde (pan-membrane-directed) vesicle trajectories. This regulatory aspect principally distinguishes the here described process of actin-dependent apical vesicle delivery from the formin-regulated actin-dependent apical vesicle delivery that serves to expand but not to position the inter- and intracellular apical/lumenal membrane in *Drosophila* tubular epithelia and requires track-dependent MyoV-driven vesicle movement[41] (such tracks also function in pancreatic secretion[42]; and from the Arp2/3-dependent apical trafficking of Delta into apical microvilli of *Drosophila* sensory organ precursors[43]. Similar mechanisms of directed vesicle delivery may, however, constitute a late step during a process of membrane polarization that involves the asymmetric insertion of the apical domain. Moreover, the mutual interaction of endo-/vesicle membranes and actin dynamics in self-organizing networks can generate new directions for delivery routes that expand on diverse mechanism of actin-based vesicle movement. For instance, the formin-dependent movement of RAB-11 vesicles on unbranched actin filaments to the cortex of non-polarized mouse oocytes [44, 45]was subsequently shown to be modified by vesicle density to direct asymmetric spindle positioning [46]. Likewise, a combination of a subcortical actin meshwork and organelle-based actin clouds that can transform into comet tails both shuffle and direct mitochondria during cell division [47, 48]. Any vectorial (apical) actin movement, powered by branched-chain actin dynamics, should therefore be able to confer asymmetric directionality to those vesicles that might attach to it via their previously identified apical polarity cues [5].

Our colocalization studies did not suggest (albeit not exclude) that actin comets or plumes are generated on specific vesicle populations during *de novo* polarized membrane biogenesis in the expanding *C. elegans* intestine. By confocal imaging using LifeAct::GFP, the here-described actin structure more closely resembles the actin mesh proposed to maintain microtubule organization in the *Drosophila* oocyte, a structure considered to be static and isotropic and one that derives its power from straight-chain (formin), not branched-chain (bcAM), regulated filament assembly [49]. In further contrast , the here described actin structure is self-organizing, directional, and dynamic. Intriguingly, it appears to require previously unidentified, almost identical *C. elegans* intestinal actin isoforms, isoforms chiefly expressed in muscle, yet required for intestinal polarity. The combinatorial 3'UTR approach used for their analysis demonstrates that their function in polarity is not redundant and not simply a matter of quantity. We suggest that the identified isoforms contribute structural specificity to this novel actin structure (the specific requirement of isoforms is an emerging theme in the higher-resolution dissection of cytoskeletal assemblies; compare [50]. Most remarkably, however, photomodulation using LifeAct demonstrates that this filamentous actin structure, driven by bcAMs, asymmetrically moves or retracts towards the apical domain.

Filamentous actin assemblies are best known for their structural-mechanical maintenance and force-mechanical biochemical or physical signal transduction functions, but also have the ability to self-organize and generate flow. Actin treadmilling, executed with the help of F-actin assembly modifiers (branched and straight chain), classically illuminates actin's self-organizing potential. It operates in systems as ancient as protozoans (ActA-dependent forward propulsion of listeria [46, 51] and informs these protozoan's ability to exploit actin networks of multicellular organisms, including those of the *C. elegans* intestine [52]. A combination of treadmilling and regulated Arp2/3 and capping protein dependent actin mesh assembly drives lamellipodial protrusions during directed cell migration, with speeds saturable at high actin concentrations[53]. However, the translocation of photoactivated or converted LifeAct fluorophores observed here suggests that, rather than repeat cycles of filament assembly and disassembly that generate the impression of movement, it is a polymeric F-actin structure that actually moves during polarity establishment in the *C. elegans* intestine.

A defining characteristic of this structure is its contiguity with the cell cortex and its basolateral-to-apical translocation along the cortex during polarity establishment. Cortical actin modeling, typically driven by actomyosin activity and involved in the polarity of non-epithelial cells, has recently also been implicated in the apical polarity of epithelial cells (see above and [54]. Moreover, it is a classical polarity cue in the *C. elegans* zygote. Activated by sperm entry, actin dynamics are subsequently self-propagating, and they include cortical dynamics as well as cytoplasmic flow, both of which operate upstream of PARs in the establishment and maintenance of zygote polarity [55]. New forms of actin filament alignment along memorized directional paths take place at the cortex[56]. However, the immediate purpose of cortex-based actin dynamics is the partitioning of the cortex itself, e.g., the polarized distribution of PARs, not the specification of intracellular events such as polarized trafficking (also in [54]), a possibility not yet investigated in the *C. elegans* zygote (we note, however, that bcAMs direct eondosomal recycling of PAR-6 to the anterior cortex to maintain zygote polarity [57]. Cortical actomyosin networks are also thought to directly effect cortex polarization in the 8-cell mouse embryo, upstream of PARs [58]. Set into motion by an extrinsic, signal-dependent orientation of the cortical actin network (sperm entry in the *C.elegans* zygote, PLC-dependent $PIP_2$ hydrolysis in the mammalian embryo), membrane- or cortex-based events are thus viewed as the initial step in polarity establishment, consistent with current polarity models. However, cortical actin dynamics may also complement the here proposed polarity model, e.g., via the concomitant or subsequent (with the directional delivery of apical membrane) polarized recruitment of PARs, a process that remains poorly understood. For instance, it is tempting to speculate that a cortex-based self-organizing vectorial actin assembly may also support the lateral-to-apical movement of the earliest intestinal polarity cue, PAR-3, in the *C. elegans* intestine [59]. In favor of this speculation and of the here proposed alternative polarity model, RAB-11 is already polarized in the pre-intercalation/pre-polarization embryonic *C. elegans* intestine (at the E8 stage), i.e., before PAR-3 assembles at the midline (the future apical domain), and it moves with the midbody to the future apical membrane[60].

Beyond cortical actin dynamics, cytoplasmic flow dynamics of actin networks are increasingly appreciated as another mechanism that can confer directionality to cellular processes. For example, cytoplasmic flow dynamics have taken center stage over canonical leading-edge movement in directed cell migration, where gradients of cytoplasmic actin network compression and destruction, regulated by myosin and cofilin, have been identified [61]. Directional cytoplasmic bulk actin dynamics induces phase segregation of ooplasm and yolk granules in zebrafish oocytes that induces comet assembly at the yolk granules which pushes them into the opposite direction [62]. The full molecular, structural, and mechanical characterization of the here described vectorial polymeric F-actin structure should provide further insight into the still enigmatic process of polarity establishment and maintenance.

**References.**

14. Stinchcomb, D.T., Shaw, J.E., Carr, S.H. & Hirsh, D. Extrachromosomal DNA transformation of Caenorhabditis elegans. *Mol Cell Biol* **5**, 3484-3496 (1985).
15. Mello, C.C., Kramer, J.M., Stinchcomb, D. & Ambros, V. Efficient gene transfer in C.elegans: extrachromosomal maintenance and integration of transforming sequences. *EMBO J* **10**, 3959-3970 (1991).
16. Gobel, V., Barrett, P.L., Hall, D.H. & Fleming, J.T. Lumen morphogenesis in C. elegans requires the membrane-cytoskeleton linker erm-1. *Dev Cell* **6**, 865-873 (2004).
17. Khan, L.A. *et al.* Intracellular lumen extension requires ERM-1-dependent apical membrane expansion and AQP-8-mediated flux. *Nature cell biology* **15**, 143-156 (2013).
18. Zhang, H. *et al.* Clathrin and AP-1 regulate apical polarity and lumen formation during C. elegans tubulogenesis. *Development* **139**, 2071-2083 (2012).
19. Ramalho, J.J. *et al.* C-terminal phosphorylation modulates ERM-1 localization and dynamics to control cortical actin organization and support lumen formation during Caenorhabditis elegans development. *Development* **147** (2020).
20. Mariol, M.C., Walter, L., Bellemin, S. & Gieseler, K. A rapid protocol for integrating extrachromosomal arrays with high transmission rate into the C. elegans genome. *J Vis Exp*, e50773 (2013).
21. Griffin, E.E., Odde, D.J. & Seydoux, G. Regulation of the MEX-5 gradient by a spatially segregated kinase/phosphatase cycle. *Cell* **146**, 955-968 (2011).
22. Patterson, G.H. & Lippincott-Schwartz, J. A photoactivatable GFP for selective photolabeling of proteins and cells. *Science* **297**, 1873-1877 (2002).
23. Mijalkovic, J., Prevo, B., Oswald, F., Mangeol, P. & Peterman, E.J. Ensemble and single-molecule dynamics of IFT dynein in Caenorhabditis elegans cilia. *Nat Commun* **8**, 14591 (2017).
24. Schneider, C.A., Rasband, W.S. & Eliceiri, K.W. NIH Image to ImageJ: 25 years of image analysis. *Nat Methods* **9**, 671-675 (2012).
25. He, J. *et al.* Prevalent presence of periodic actin-spectrin-based membrane skeleton in a broad range of neuronal cell types and animal species. *Proc Natl Acad Sci U S A* **113**, 6029-6034 (2016).
26. Candelli, A., Wuite, G.J. & Peterman, E.J. Combining optical trapping, fluorescence microscopy and micro-fluidics for single molecule studies of DNA-protein interactions. *Phys Chem Chem Phys* **13**, 7263-7272 (2011).
27. Erb, I. *et al.* Use of ChIP-Seq data for the design of a multiple promoter-alignment method. *Nucleic Acids Res* **40**, e52 (2012).
28. Labouesse, M. Epithelial junctions and attachments. *WormBook*, 1-21 (2006).
29. Madhu, B., Salazar, A. & Gumienny, T. Caenorhabditis elegans egg-laying and brood-size changes upon exposure to Serratia marcescens and Staphylococcus epidermidis are independent of DBL-1 signaling. *MicroPubl Biol* **2019** (2019).
30. Di Tommaso, P. *et al.* T-Coffee: a web server for the multiple sequence alignment of protein and RNA sequences using structural information and homology extension. *Nucleic Acids Res* **39**, W13-17 (2011).
31. Ono, K., Parast, M., Alberico, C., Benian, G.M. & Ono, S. Specific requirement for two ADF/cofilin isoforms in distinct actin-dependent processes in Caenorhabditis elegans. *J Cell Sci* **116**, 2073-2085 (2003).
32. Asan, A., Raiders, S.A. & Priess, J.R. Morphogenesis of the C. elegans Intestine Involves Axon Guidance Genes. *PLoS Genet* **12**, e1005950 (2016).

33. Cao, J. *et al.* Comprehensive single-cell transcriptional profiling of a multicellular organism. *Science* **357**, 661-667 (2017).
34. MacQueen, A.J. *et al.* ACT-5 is an essential Caenorhabditis elegans actin required for intestinal microvilli formation. *Mol Biol Cell* **16**, 3247-3259 (2005).
35. Landel, C.P., Krause, M., Waterston, R.H. & Hirsh, D. DNA rearrangements of the actin gene cluster in Caenorhabditis elegans accompany reversion of three muscle mutants. *J Mol Biol* **180**, 497-513 (1984).
36. Waterston, R.H., Hirsh, D. & Lane, T.R. Dominant mutations affecting muscle structure in Caenorhabditis elegans that map near the actin gene cluster. *J Mol Biol* **180**, 473-496 (1984).
37. Avery, L., Bargmann, C.I. & Horvitz, H.R. The Caenorhabditis elegans unc-31 gene affects multiple nervous system-controlled functions. *Genetics* **134**, 455-464 (1993).
38. Avery, L. The genetics of feeding in Caenorhabditis elegans. *Genetics* **133**, 897-917 (1993).
39. Avery, L. Motor neuron M3 controls pharyngeal muscle relaxation timing in Caenorhabditis elegans. *J Exp Biol* **175**, 283-297 (1993).
40. Stone, S. & Shaw, J.E. A Caenorhabditis elegans act-4::lacZ fusion: use as a transformation marker and analysis of tissue-specific expression. *Gene* **131**, 167-173 (1993).
41. Massarwa, R., Schejter, E.D. & Shilo, B.Z. Apical secretion in epithelial tubes of the Drosophila embryo is directed by the Formin-family protein Diaphanous. *Dev Cell* **16**, 877-888 (2009).
42. Geron, E., Schejter, E.D. & Shilo, B.Z. Directing exocrine secretory vesicles to the apical membrane by actin cables generated by the formin mDia1. *Proc Natl Acad Sci U S A* **110**, 10652-10657 (2013).
43. Rajan, A., Tien, A.C., Haueter, C.M., Schulze, K.L. & Bellen, H.J. The Arp2/3 complex and WASp are required for apical trafficking of Delta into microvilli during cell fate specification of sensory organ precursors. *Nat Cell Biol* **11**, 815-824 (2009).
44. Schuh, M. An actin-dependent mechanism for long-range vesicle transport. *Nat Cell Biol* **13**, 1431-1436 (2011).
45. Pfender, S., Kuznetsov, V., Pleiser, S., Kerkhoff, E. & Schuh, M. Spire-type actin nucleators cooperate with Formin-2 to drive asymmetric oocyte division. *Curr Biol* **21**, 955-960 (2011).
46. Holubcova, Z., Howard, G. & Schuh, M. Vesicles modulate an actin network for asymmetric spindle positioning. *Nat Cell Biol* **15**, 937-947 (2013).
47. Moore, A.S. & Holzbaur, E.L.F. Actin mixes up mitochondria for inheritance. *Nature* (2021).
48. Moore, A.S. *et al.* Actin cables and comet tails organize mitochondrial networks in mitosis. *Nature* **591**, 659-664 (2021).
49. Dahlgaard, K., Raposo, A.A., Niccoli, T. & St Johnston, D. Capu and Spire assemble a cytoplasmic actin mesh that maintains microtubule organization in the Drosophila oocyte. *Dev Cell* **13**, 539-553 (2007).
50. Khan, L.A. *et al.* A tensile trilayered cytoskeletal endotube drives capillary-like lumenogenesis. *J Cell Biol* **218**, 2403-2424 (2019).

51. Kocks, C., Hellio, R., Gounon, P., Ohayon, H. & Cossart, P. Polarized distribution of Listeria monocytogenes surface protein ActA at the site of directional actin assembly. *J Cell Sci* **105 ( Pt 3)**, 699-710 (1993).
52. Estes, K.A., Szumowski, S.C. & Troemel, E.R. Non-lytic, actin-based exit of intracellular parasites from C. elegans intestinal cells. *PLoS Pathog* **7**, e1002227 (2011).
53. Hu, L. & Papoian, G.A. Mechano-chemical feedbacks regulate actin mesh growth in lamellipodial protrusions. *Biophys J* **98**, 1375-1384 (2010).
54. Zihni, C. *et al.* An apical MRCK-driven morphogenetic pathway controls epithelial polarity. *Nat Cell Biol* **19**, 1049-1060 (2017).
55. Munro, E., Nance, J. & Priess, J.R. Cortical flows powered by asymmetrical contraction transport PAR proteins to establish and maintain anterior-posterior polarity in the early C. elegans embryo. *Dev Cell* **7**, 413-424 (2004).
56. Li, Y. & Munro, E. Filament-guided filament assembly provides structural memory of filament alignment during cytokinesis. *Dev Cell* **56**, 2486-2500 e2486 (2021).
57. Shivas, J.M. & Skop, A.R. Arp2/3 mediates early endosome dynamics necessary for the maintenance of PAR asymmetry in Caenorhabditis elegans. *Mol Biol Cell* **23**, 1917-1927 (2012).
58. Zhu, M., Leung, C.Y., Shahbazi, M.N. & Zernicka-Goetz, M. Actomyosin polarisation through PLC-PKC triggers symmetry breaking of the mouse embryo. *Nat Commun* **8**, 921 (2017).
59. Melissa A. Pickett, M.D.S., Victor F. Naturale, Deniz Akpinaroglu, Joo Lee, Kang Shen, Jessica L. Feldman Separable mechanisms drive local and global polarity establishment in the C. elegans intestinal epithelium. *bioRxiv* (2021).
60. Bai, X. *et al.* Aurora B functions at the apical surface after specialized cytokinesis during morphogenesis in C. elegans. *Development* **147** (2020).
61. Yolland, L. *et al.* Persistent and polarized global actin flow is essential for directionality during cell migration. *Nat Cell Biol* **21**, 1370-1381 (2019).
62. Shamipour, S. *et al.* Bulk Actin Dynamics Drive Phase Segregation in Zebrafish Oocytes. *Cell* **177**, 1463-1479 e1418 (2019).
